# Host evolutionary history predicts virus prevalence across bumblebee species

**DOI:** 10.1101/498717

**Authors:** David J. Pascall, Matthew C. Tinsley, Darren J. Obbard, Lena Wilfert

**Author notes:** Institute of Biodiversity, Animal Health and Comparative Medicine, Graham Kerr Building, University of Glasgow, Glasgow, UK. Institute of Evolutionary Ecology and Conservation Genomics, University of Ulm, Albert-Einstein-Allee 11, 89069 Ulm, Germany (Lena Wilfert).

## Abstract

Why a pathogen associates with one host but not another is one of the most important questions in disease ecology. Here we use transcriptome sequencing of wild-caught bumblebees from 13 species to describe their natural viruses, and to quantify the impact of evolutionary history on the realised associations between viruses and their pollinator hosts. We present 37 novel virus sequences representing at least 30 different viruses associated with bumblebees. We verified 17 of them by PCR and estimate their prevalence across species in the wild. Through small RNA sequencing, we demonstrate that at least 10 of these viruses form active infections in wild individuals. Using a phylogenetic mixed model approach, we show that the evolutionary history of the host shapes the current distribution of virus/bumblebee associations. Specifically, we find that related hosts share viral assemblages, viruses differ in their prevalence averaged across hosts and the prevalence of infection in individual virus-host pairings depends on precise characteristics of that pairing.

**Author’s Summary:** Despite the importance of disease in the regulation of animal populations, our understanding of the distribution of pathogen burden across wild communities remains in its infancy. In this study, we investigated the distribution of viruses across natural populations of 13 different bumblebee species in Scotland. We first searched for viruses using a metatranscriptomic approach, finding at least 30 new viruses of bumblebees, and assayed a subset of them for their presence and absence in different host species. Then, in the first application of these methods to an animal-virus system, we used co-phylogenetic mixed models to investigate the factors that lead to species being having different prevalences for a subset of these viruses. While much of the variation in the prevalence of the viruses can be explained by the idiosyncrasies of individual bumblebee-virus pairings, there is a phylogenetic signal with related bumblebee species being infected at similar frequencies by the same sets of viruses. Consistent with previous work, our study indicates that, while in general the interaction between a host and a virus may be unpredictable, closely related species are more likely to exhibit similar patterns of infection.

## Introduction

Pathogens that naturally infect more than one host species have a particularly high risk of disease emergence [1]. One especially important group of pathogens are the viruses, whose ubiquity leads them to have a disproportionate role in the regulation of natural populations [2]. Viruses are relevant in populations that humans manage for economic and conservation reasons, such as bumblebees, which are both in decline [3] and important providers of ecosystem services [4].

Bumblebees, genus *Bombus*, are a primitively eusocial group of important wild pollinators. Many bumblebee species have experienced population declines, linked to biotic and abiotic stressors such as habitat degradation, pesticide use and shared infectious diseases for example caused by viral pathogens [5]. In contrast to honeybee viruses, which have been intensively studied, and have in many cases been found to represent multihost pathogens (see Manley et al (2015) [6] and the references within), bumblebee-specific viruses are comparatively poorly studied, and it is unknown how widely they are shared between host species.

For a species to be a multihost pathogen, some degree of opportunity for cross-species transmission must exist. Our definition of multihost pathogens follows that of Fenton et al. [7]. As such, multihost pathogens are defined to include two conceptually distinct groups: ‘facultative multihost pathogens’ that are able to maintain transmission chains in multiple host species (i.e. *R*_*0*_>=1 in multiple host species) and ‘obligate multihost pathogens’, which rely on sufficiently high rates of cross-species transmission to offset unsustainable transmission within individual host species (i.e. 0<*R*_*0*_<1 within host species, *R*_*0*_>=1 overall). In addition, pathogens that maintain transmission in a single host (*R*_*0*_>=1) but experience regular spillover (with or without the expectation of onward transmission: 0<= *R*_*0*_<1) are included as being effectively multihost pathogens within our definition. *R*_*0*_ is defined as the expected number of secondary infections caused by a single typical infected individual in an entirely naïve host population [8]. We define cross-species transmission as the movement of a multihost pathogen between host species within its host range. This contrasts with host shifting, which we define as a transmission event to a new host species, leading to a change in host range. However there is necessarily some unavoidable ambiguity between cross-species transmission and host shifting in the case of pathogens that exhibit rare spillover events.

The opportunity for cross-species transmission, which explains the large number of viruses originally detected in honeybees that are present in bumblebees, may be created by niche overlap in foraging [9]. Bumblebee nests are provisioned by foraging workers who gather pollen and nectar from flowers in the surrounding area. Considerable interspecific differences in plant species utilization by foragers of different species are commonly observed [10-13], but this is not a universal phenomenon [14], and the degree of overlap may depend on the diversity of flowers currently in bloom. In bumblebees, flower choice of foragers is correlated with species tongue length [12,13], which implicitly incorporates shared behavioural characteristics between closely related bumblebee species as there is phylogenetic correlation between tongue length and relatedness [15]. Different species of bumblebee also exhibit incomplete temporal separation throughout the year, causing some degree of partitioning in niche space even when they are spatially sympatric [16]. This ecology leads to a complex interaction network between bumblebee species, as well as sympatric honeybees, which may structure cross-species transmission.

The prevalence of pathogens, including viruses, across host species, such as bumblebees, is structured on two levels. First, a virus may be present or entirely absent in a potential host species. Second, other factors may then influence how prevalent a pathogen is within that species. At the presence/absence level, a complete lack of infection in nature can occur in three ways: 1) a host and virus may exist in allopatry or in completely non-interacting ecological niches, preventing transmission irrespective of the host’s susceptibility; 2) a physiological or molecular mismatch (including immunity) between a host and virus can prevent infection; and 3) environmental conditions may be such that transmission cannot occur between two sympatric species. None of these mechanisms represent an immutable barrier, and all represent ends of a continuum, where lesser forms simply reduce transmission. Spatially or ecologically separated hosts and parasites may come into contact through migrations or human facilitated invasions, allowing new associations to emerge. For example, the arrival of *Plasmodium relictum* to the Hawaiian islands led to avian population declines and contributed to extinctions in the naturally susceptible but naïve populations [17]. Incompatibility can break down if evolution in the pathogen or host removes the physiological or molecular barriers to infection, as shown when Canine parvovirus type 2 emerged from Feline panleukopenia virus after gaining the ability to bind to canine transferrin receptors [18].

For virus-host associations where infection can and does occur, quantitative differences in infection risk between species can be driven by ecological variation in transmission rates. These differences can be driven by, for example, the propensity for group living [19], population densities [20], the biodiversity of the community [21] and host avoidance behaviours [22]. Variation in infection risk among host species can also be driven by physiological and molecular factors, with hosts having varying suitability for the replication of a given parasite. In the extreme case, a host species may exhibit condition-dependent susceptibility; where infection can only occur when the immune system is suppressed, either directly, through an immunosuppressant disease or chemical agent, or indirectly, through trade-offs in resource allocation brought about by malnutrition [23]. Both behavioural and ecological factors, leading to differences in contact rate, and physiological factors, leading to differences in infection probability on contact, may be phylogenetically correlated [15,24].

### Box 1 – Definition of Terms

Co-phylogenetic generalized linear mixed models that incorporate phylogenetic variance from multiple clades [25,26] have been used relatively rarely, and a biological interpretation of the model terms may not be immediately familiar. In the host-parasite context, this approach can be used to model how the probability of infection is predicted by both host and parasite species, allowing for covariance induced by the relationships within each group, and the interactions between these model terms. This can be considered either at the species-wide level (i.e. the probability that infection will occur at all in a given host/parasite pairing), or at the level of individuals within species (i.e. infection prevalence). Here we provide verbal descriptions of how the terms can be interpreted, as well as references to a figure in Hadfield et al. (2014) [25] where each of these effects is illustrated graphically:

#### Phylogenetic Effect

Variation in the mean value of a trait among species that is explained by phylogenetic divergence. For example, more closely-related hosts might be more similar in susceptibility to viral infection (display higher viral prevalence), irrespective of virus species. Equivalently, more closely-related viruses might be more similar in infectivity, irrespective of host species (Figs 1a and b in Hadfield et al. (2014) [25]).

**Fig 1.**
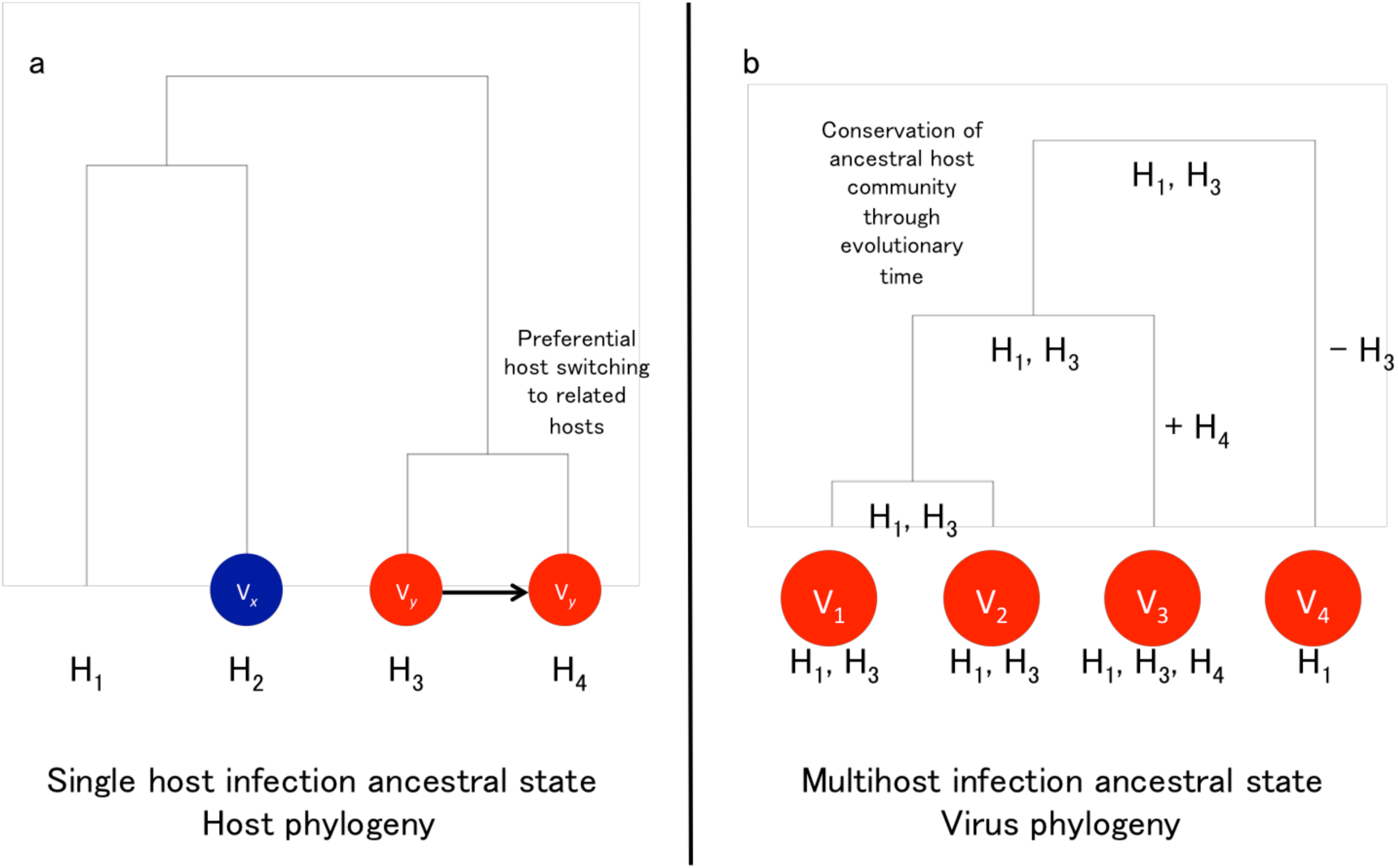
Mechanisms for the generation of novel multihost viruses. The generation of novel multihost viruses through host shifting (1a) leads to a ‘host evolutionary interaction’ effect (Box1), as the consistent switching of viruses (V) to hosts (H) closely related to their ancestral host will lead to related hosts having correlated viral assemblages. The generation of novel multihost viruses through speciation (1b) can lead to a ‘virus evolutionary interaction’ effect (Box 1) through the inheritance of the ancestral host range, leading to the daughter virus species having correlated host assemblages.

**Fig 2.**
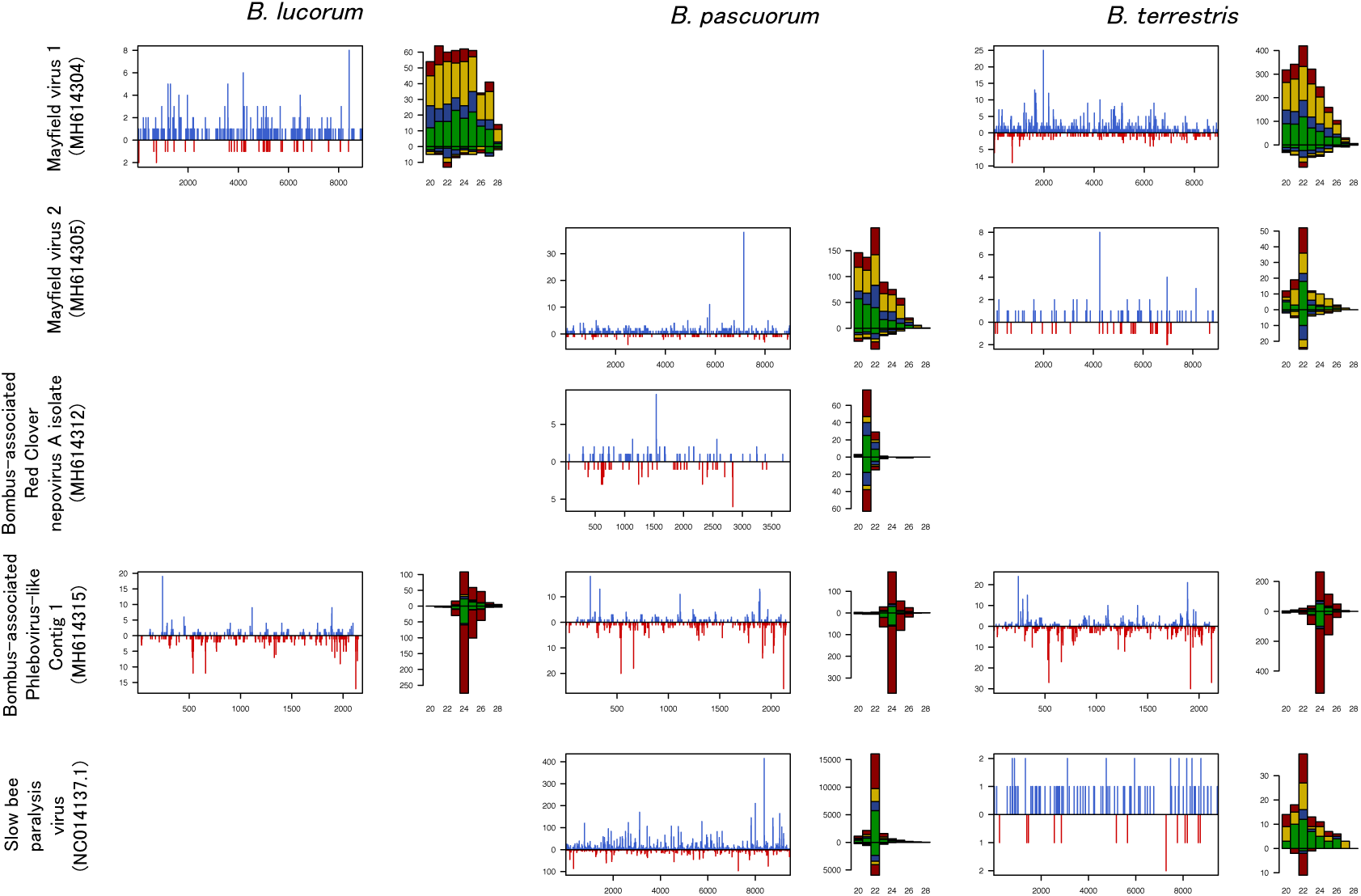
The mapping of small RNA reads to Mayfield virus 1, Mayfield virus 2, a contig similar to Tomato black ring virus, a contig similar to a phlebovirus glycoprotein and Slow bee paralysis virus. Blue lines represent reads mapping to the positive-sense strand at that genomic position, red lines represent reads mapping to the negative sense strand. The histogram of read size spectra shows the count of reads of each length mapping in the positive (above) and negative (below) directions. The colouring of each bar shows the counts of the reads beginning with each 5’ base (red-U, blue-C, green-A, yellow-G).

#### Species Effect

Variation in the mean value of a trait among species that is not explained by a *Phylogenetic Effect*. For example, much of the variation in prevalence among viral species (irrespective of host), may not be explained by the virus phylogeny but instead depend on lineage-specific viral traits (Figs 1g and f in Hadfield et al. (2014) [25]).

#### Non-phylogenetic Interaction

The interaction term between host and parasite *Species Effects*, such that variation in the mean value of a trait within particular host/parasite pairings depends the specifics of the host and parasite involved in a way not affected by their evolutionary divergence. For example, variation in prevalence between particular pairings that is caused by the interaction between lineage-specific host and viral traits (Fig 1h in Hadfield et al. (2014) [25]).

#### Coevolutionary Interaction

The interaction term between host and parasite *Phylogenetic Effects*, such that variation in the mean value of a trait within particular host/parasite pairings depends on the evolutionary divergence among species in both host and parasite clades. For example, if the prevalence of infection is more similar among pairings of closely-related hosts and closely-related parasites than would be expected from the host and parasite phylogenies and species-means alone. (Fig 1e in Hadfield et al. (2014) [25]).

#### Evolutionary Interaction

The interaction term between the host (or parasite) *Phylogenetic Effect* and the partners’ *Species Effect*, such that the variation in the mean value of a trait within particular host/parasite pairings depends on the evolutionary divergence among hosts (or parasites) and on the identity of particular partner species, but is not predicted by the evolutionary divergence between partners’ species. For example, if the similarity in viral prevalence for one virus species is strongly predicted by the evolutionary divergence among hosts, but a completely different relationship (unrelated to the evolutionary divergence among viruses) is seen for other virus species. (Fig 1c and d in Hadfield et al. (2014) [25]).

A new multihost virus can arise in two ways, either through a virus species gaining a new host [27] (Fig 1a), or through a speciation event in a multihost virus (i.e. giving rise to two sibling multihost virus species; Fig 1b). When novel multihost viruses are generated through host shifting (Fig 1a), a ‘host evolutionary interaction’ effect (see Box 1) can result, as the consistent switching of viruses (V) to hosts (H) closely related to their ancestral host will lead to related hosts having correlated viral assemblages. When novel multihost viruses arise through speciation, i.e. if the ability to infect multiple hosts is an ancestral trait (Fig 1b), a ‘virus evolutionary interaction’ effect can result (see Box 1) through the inheritance of the ancestral host range, leading to the daughter virus species having correlated host assemblages. These effects can also be generated in ecological time through mechanisms that lead to biased cross-species transmission.

Here we test for a role of evolutionary history in shaping the current host/virus assemblage using species from an ecologically and economically important group, the bumblebees. We cataloged the virome of wild-caught bumblebees from across Scotland by RNAseq, finding at least 30 new viruses. We then tested multiple bumblebee species for a subset of these novel viruses and three previously reported honeybee viruses: Slow bee paralysis virus [28], Acute bee paralysis virus [29] and Hubei partiti-like virus 34 [30,31]. We analysed virus prevalence using co-phylogenetic models to determine the presence or absence and relative strengths of the phylogenetic signals that are expected to shape the host/virus assemblage in this system, and performed tests to attempt to determine the mechanisms driving this.

## Methods

### Sampling strategy

A total of 926 individual bumblebees of 13 species were collected on the wing from nine sites across Scotland in July and August of 2009 and 2011, and frozen in liquid nitrogen or at −80°C. In 2009, we sampled the Ochil Hills, Glenmore, Dalwhinnie, Stirling, Iona, Staffa, and the Pentlands, and in 2011 we sampled Edinburgh and Gorebridge (Supp Table 1). The cryptic species complex of *Bombus terrestris, Bombus lucorum, Bombus cryptarum* and *Bombus magnus* was resolved using RFLP analysis following Murray et al. (2008) [32]. All individuals were bisected longitudinally prior to RNA extraction. One half of each bumblebee was used in pooled RNA extractions of 2-11 individuals per species (median 10; Supp Table 2). Two of these pools (‘DIV’ and ‘P11’) were included in the RNAseq, but excluded from prevalence testing. The groups of bumblebees were ground in liquid nitrogen and added to TRIzol reagent (Life Technologies) for RNA extractions, following the manufacturer’s standard protocol. The RNA concentrations in the pooled samples were equalized to approximately 200 ng/ul/individual based on Nanodrop measurements.

### RNA Sequencing and Bioinformatics

The RNA was combined by species for *B. terrestris* (239 individuals), *Bombus pascuorum* (212 individuals), *B. lucorum* (182 individuals) and other *Bombus* (293 individuals) into four large RNA pools. These large pools were sequenced using the Illumina HiSeq platform with 100bp paired end reads (Beijing Genomics Institute) after poly-A selection. This excludes ribosomal and bacterial RNA, and will enrich for mRNAs and those RNA viruses that have polyadenylated genomes or products. The single-species bumblebee pools were subsequently re-sequenced following duplex specific nuclease normalization, to reduce rRNA representation, and enrich for rare transcripts while retaining non-polyadenylated viruses and products. The small RNAs of the same RNA pools of *B. terrestris, B. lucorum* and *B. pascuorum* were also sequenced to test for the replication of viruses identified via the transcriptome sequencing.

For each pool, paired end RNAseq data were initially mapped to the published *Bombus terrestris* and *B. impatiens* genomes using bowtie2 [33] to reduce the representation of conserved bumblebee sequences. Read pairs that did not map concordantly, including divergent bumblebee sequences and other associated microbiota, were assembled *de novo* using Trinity 2.2.0 [34] as paired end libraries, following automated trimming (‘--trimmomatic’) and digital read normalisation (‘--normalize_reads’). Where two RNAseq libraries (Poly-A and DSN) had been sequenced, these were combined for assembly.

To identify putative viruses, all long open reading frames from each contig were identified and concatenated to provide a ‘bait’ sequence for similarity searches using Diamond [35] and BLASTp [36]. Contigs shorter than 500 base pairs were discarded. These contig translations were used to search against a Diamond database comprising all of the virus protein sequences available in NCBI database ‘nr’, and all of the Dipteran, Hymenopteran, Nematode, Fungal, Protist, and prokaryotic proteins available in NCBI database ‘refseq_protein’ (mode ‘blastp’; e-value 0.001; maximum of one match). Matches to phage and short matches to large DNA viruses were excluded. Remaining contigs were manually curated to identify and annotate high-confidence virus-like sequences. To quantify approximate fold-coverage, and to assess viRNA properties, the raw RNAseq and trimmed small RNA reads were mapped against the putative viral contigs using bowtie2’s ‘--very-sensitive’ setting and retaining only the top map [33], from this we recorded the number of mapped reads per kilobase of transcript per million mapped reads. We considered viruses to show strong evidence of replication in the host if they had at least 50 mapping siRNA reads with a size distribution sharply peaked at 22nt (viRNAs are generated from replicating viruses by Dcr2). Following Fauquet and Stanley (2005) [37], we designated contigs exhibiting less than 90% nucleotide identity as separate viruses and those exhibiting greater than or equal to 90% identity as strains of known viruses.

### PCR Validation and Testing

A subset of contigs were chosen for manual validation. All chosen contigs met both of the following conditions: the presence of mapping reads in the bumblebee small RNAs (for the *B. terrestris, B. pascuorum* and *B. lucorum* pools; not a condition for the mixed *Bombus* pool) or the transcriptomic RNAs (for the other *Bombus* pool where small RNAs were not generated), and the closest blast match being viral RNA-dependent RNA-polymerase. Internal primers for these contigs were generated using primer3 [38] and amplification of the target was verified via Sanger sequencing. See Supp Table 3 for PCR conditions and primer sequences. Mayfield virus 1 and 2 were Sanger sequence validated over the entirety of the contig. The Loch Morlich and River Liunaeg virus sequences were generated by the connection of several disjoint contigs by Sanger sequencing. Black Hill virus was excluded from further analysis as it was found that that the PCR reaction amplified a host sequence that could not be visually differentiated from the virus product.

### Phylogenetic Inference

Following Cameron et al. (2007) [39], we inferred the bumblebee phylogeny using cytochrome oxidase I, elongation factor 1-alpha, opsin, phosphoenolpyruvate carboxykinase, 16S and arginine kinase genes. To break up long branches and allow dating, additional species not sampled in the field were added from public databases (see Supp Table 4 for genbank accession numbers and species included). The DNA sequences were aligned with MAFFT using the L-INS-i setting [40,41]. The 6 gene alignments were then used to generate the phylogeny in BEAST v2.4.5 [42], treating each file as a separate partition, using bModelTest [43] with the ‘transitionTransversion split’ setting and a calibrated Yule tree prior [44]. An uncorrelated lognormal relaxed clock was fitted to each partition, with exponential (λ=1) priors placed over the mean rate and the default gamma (α=0.5396, β=0.3819) priors being placed over the standard deviation [45]. The bumblebees were constrained to be monophyletic, with the honeybee, *Apis mellifera*, as an outgroup. A gamma (α=74.85889, β=0.4366812) distributed divergence time prior was placed over the tMRCA of the *Bombus* clade, with parameters optimised to match the 2.5^th^ and 97.5^th^ percentiles of the posterior distribution of ages previously estimated by Hines (2008) [46]. Four separate runs of the MCMC were performed for 100,000,000 steps from random starting trees, with the first 50,000,000 steps being discarded as burn in. Convergence of the posterior among runs was assessed in Tracer v1.7 [47]. The posterior sample was thinned to 1000 trees.

For the virus phylogeny, amino acid sequences were inferred based on the translated ORFs for regions predicted to contain RdRp motifs using the GenomeNet MOTIF search function [48] against the Pfam database [49], with an expectation cut-off of 0.00001. If a virus had no annotated motifs, the canonical GDD RdRp amino acid motif [50] was identified manually. Additional virus species (Supp Table 5) were added to the phylogeny to anchor species with short generated contigs, and to break up long branches. Given the long evolutionary distance between the viruses, PROMALS3D [51] was used to align viral sequences. The alignments were trimmed to the first conserved secondary structural element at both ends as predicted by PROMALS3D with the 0.95 conservation metric. Two of the novel viruses (Agassiz Rock virus and Cnoc Mor virus) were not included in this phylogeny because the section of the RdRp gene required fell outside the available contig. Given that it is unclear whether there was a universal common ancestor of all RNA viruses [52], we aligned the sequences and generated the phylogeny twice, with and without the negative sense RNA viruses (Supp Table 5). The trees serve purely to quantify expected variance (under a Brownian motion model of evolution) between closely related viruses. The deep splits in the phylogeny are poorly resolved with RdRp data [53], due to the fast evolutionary rates of RNA viruses, the considerable time since divergence and permutations in the RdRp sequence [54]. However, this should not overly bias the conclusions as beyond a certain evolutionary distance, the viruses would be expected to become essentially uncorrelated when averaged across the posterior (Supp Table 6 for realised correlations).

Phylogenetic models used the BLOSUM62 rate matrix [55] with gamma distributed rate variation using 4 gamma categories, an uncorrelated lognormal relaxed clock [45] and a Yule tree prior. A CTMC rate reference prior [56] was placed over the clock mean and an exponential (λ=1) prior was placed over the standard deviation. The alpha parameter of the gamma distributed rate variation was given an exponential (λ=1) prior. Absolute dating of viral trees is difficult due to the inconsistency in estimated ages provided by estimated clock rates and known orthologous insertions between sister host species [57], but is not essential for our analysis, which depends only on relative branch lengths. Nevertheless, we chose to use orthologous insertions (related historical viruses stably integrated into the genomes of species with better known divergence dates) to provide approximate dates for our tree. To account for the estimated ages of RNA viral families [58], we set a uniform lognormal prior with an offset of 97 Mya, a mean of 500 Mya and a logged standard deviation of 0.5 on the age of the root of the tree including the negative sense RNA viruses and a lognormal prior with an offset of 76 Mya, a mean of 500 Mya and a logged standard deviation of 0.5 on the age of the tree excluding them. Two partitiviruses (Rosellinia necatrix partitivirus 2 and Raphanus sativus cryptic virus 1) known to have a common ancestor older than 10 Mya [59] were included for dating purposes. We placed a diffuse lognormal prior with an offset of 10 Mya, a mean of 30 Mya and a logged standard deviation of 0.5, on the age of the MRCA of these species. Both models were run over 10 separate chains for 50,000,000 generations on a cluster in BEAST v1.8.4 [60], with 25,000,000 generations being discarded as burn-in. Convergence of the posterior was assessed in Tracer v1.7 [47]. The posterior distributions were combined and thinned to 1000 trees.

### Prevalence Estimation

Maximum likelihood prevalence and 2-log-likelihood confidence intervals were estimated for each host/virus combination with more than one pool using the code from Webster et al. (2015) [61]. As the samples were small pooled groups of individuals, such that a PCR ‘positive’ represents one or more infections, we modelled the prevalence using a “pooled binomial” likelihood [62–64]. This approach requires that the underlying prevalence of a virus is the same in all pools, which is unlikely for bumblebees sampled from different locations. Estimates should therefore be treated with caution.

### Co-phylogenetic Mixed Model Analysis

To test for the evolutionary effects on association, the presence/absence data and the phylogenetic trees were analysed using a co-phylogenetic mixed model [25] implemented in Stan [65]. Our model is explicitly focused at the individual-level, and the model’s predictions represent the predicted probability of infection within an individual of the species. This is in contrast with Hadfield et al.’s original implementation where the focus was at the species- or population-level and the model was estimating the probability that the parasite would be found in the species or population at all. In all cases, the presented models showed no divergences, acceptable Rhat and E-BFMI values and effective sample sizes of over 200.

We fitted host and virus phylogenetic effects, which measure the extent to which variation in prevalence is clustered on the host and viral phylogenies respectively. We also fitted host and viral evolutionary interaction effects, which measure the extent to which related species have similar probabilities of infection in the sets of their interaction partners. The final phylogenetic term fitted was a coevolutionary interaction, which measures the extent to which related hosts are infected to similar degrees by related viruses (see Box 1).

In addition to the phylogenetic terms, non-phylogenetic host and virus terms, an interaction between these terms and a pool ID term were fitted. The non-phylogenetic host and virus terms measure variation that can be partitioned between host species and virus species in average infection risk that is not consistent with trait evolution by Brownian motion. The interaction term measures variation that can be partitioned between pairwise interactions between individual hosts and viruses that is not consistent with the linear sum of their individual means from the non-phylogenetic host and virus terms. The pool ID effect measures variation between pools in infection risk averaged over all the viruses tested. As the pools combined hosts by species rather than by location, so that some had individuals from multiple locations, we treated each location and each realised combination of locations as levels of a random effect, terming this the “spatial composition effect”. This describes the variation in average infection level between realised combinations of locations averaged across viruses. Model 1 included all the viruses, Model 2 excluded the negative sense RNA viruses and Model 3 fitted a pseudo-taxonomic model. In Model 3, the relationship among the viruses was represented by a polytomic viral tree with arbitrary branch lengths (with a root-to tip distance of 1 unit, and equal length between each taxonomic level) with the viruses being split first by their genomic type (+ve sense RNA, -ve sense RNA and dsRNA) implying a covariance of 0 between genome structures, followed by splitting by the putative viral clades identified by Shi et al. (2016) [31]. This was done to test for potential bias caused the by the possibility of systematic misidentification of the correct relationship between families in the estimated viral trees.

The form of the models is shown below, where *i* indexes the data points, group_*i*_ represents the level of a categorical variable that the *i*th pool belongs to, *y*_*i*_ represents the 1/0 indicator for the presence or absence of infection in the *i*th pool, *k*_*i*_ represents the number of individuals in the *i*th pool, *p*_*i*_ is the unmeasured probability of infection of a single individual in the *i*th pool, *y’*_*i*_ is the estimated value of log_*e*_(*p*_*i*_*/*(1 - *p*_*i*_)), µ is the global mean of the latent variable, ε is a normally distributed error term. All terms were fitted as random effects (i.e. estimated by partial pooling). As above, a “pooled binomial” likelihood was used [62–64].

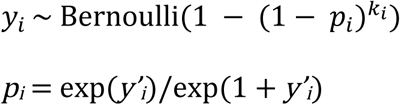

*y’*_*i*_ = µ + host_*i*_ + virus_*i*_ + interaction_*i*_ + host phylogenetic effect_*i*_ + virus phylogenetic effect_*i*_ + host evolutionary interaction_*i*_ + virus evolutionary interaction_*i*_ + coevolutionary interaction_*i*_ + pool ID;_*i*_ + species composition*i* + ε

All variance-covariance matrices were generated as described in Hadfield et al. (2014) [25], with the variance-covariance matrices scaled to correlation matrices. A standard logistic prior was placed over the global intercept on the latent scale, µ, representing a flat prior on the probability scale. An exponential (λ=1) prior was placed on each variance term in the model. In the full model with all variances being estimated, this is equivalent to a gamma (α=11, β=1) prior over the total variance, which gives a prior mean variance of 11, and an appropriate prior on the standard deviation of a variable on the logit scale. Intraclass correlations, which represent the proportion of the variance explained by each effect, were calculated on the link scale (with an addition of π^2^/3 to the denominator to account for the variance of the logistic distribution of the latent variable) from the model outputs and reported. Highest posterior density intervals were calculated by the SPIn method [66] and 90% credible intervals are reported as these are more robust to sampling in the tails of the posterior distribution [67].

The total phylogenetic variance was calculated as:

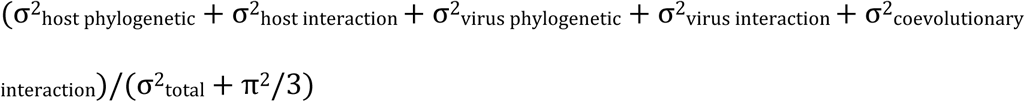

The total non-phylogenetic variance was calculated as:

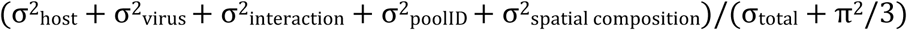

Uncertainty in the inferred phylogenies was accounted for by direct marginalisation. This dramatically increased the runtime of the model, as, given the input of *H* host phylogenies and *V* viral phylogenies from their posterior distributions, the likelihood of each datapoint has to be calculated *HV* times. As such, we included only 10 trees from each posterior, as a trade-off between runtime and accounting for uncertainty in the tree hypotheses. The marginalisation is shown below, with **y** being the total vector of presences and absences, *H* being the number of host phylogenies used, *V* being the number viral phylogenies used, θ being all the non-variance-covariance parameters in the model, Ω_*j*_ being the set of variance-covariance matrices generated by the *j*th combination of host and virus phylogenies, Ω_*HV*_ representing the set of all variance-covariance matrices being marginalised over and 𝔏 representing a likelihood.

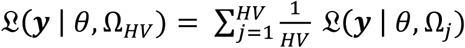

### Tongue Length Analysis

After finding that the posterior for the host evolutionary interaction was well resolved from zero, we designed a post-hoc test to attempt to detect signal for one of the obvious mechanistic explanations for this; structured transmission networks driven by evolutionarily conserved anatomical factors. We tested for an association between the tongue length differences between bumblebee species and the differences in their viral community structures, as a proxy for signal of differential transmission at flowers driven by evolutionarily conserved flower choice. Average tongue lengths for each bumblebee species except *Bombus bohemicus* and *Bombus cryptarum* were taken from Goulson et al (2005) [68]. No published tongue length could be found for *Bombus cryptarum*, so we assumed that it was identical to that of *Bombus lucorum*, a species of which it is near indistinguishable in the field. *Bombus bohemicus* was excluded from this analysis, because it is an inquiline parasite, and therefore its ecology differs from the other species in such a way that tongue length would not be expected to be correlated with the viral community distance.

In order to test for a correlation between tongue length and viral assemblage similarity, estimates of the distance in viral communities between host species are required. These were generated as follows: For each host-virus combination, the package ‘prevalence’ was used to generate posterior draws of the underlying prevalences under a Beta (1,1) prior. Then 1000 draws per species were taken from these sets of MCMC draws to generate 1000 matrices of host-virus prevalences consistent with the raw data. For each of these matrices, the distance between each species’ viral community was calculated by taking the vector of estimated prevalences for the 16 viruses of a given species as a coordinate in a 16-dimensional space then calculating the Euclidean distance between these points. The rank correlation (Kendall’s τ-b) between each pair of species’ viral community distances and their tongue length distances was then calculated, using the mantel function in the R package vegan. The point estimate presented is the median of the 1000 initial correlations accounting for the uncertainty in the underlying prevalences. The 95% confidence interval is the 2.5^th^ and 97.5^th^ percentiles distribution of estimated correlations.

## Results

RNA was extracted from 13 species of bumblebee from nine sites, to identify new viruses, assay their prevalence and their pattern of distribution across host species and to test whether the evolutionary histories of the viruses and hosts have impacted the current distribution.

### Read and Assembly Statistics

A total of 134,026,056 sequencing read pairs were generated for *Bombus lucorum*, 135,590,922 for *Bombus terrestris*, 128,670,194 for *Bombus pascuorum* and 26,838,390 for the other *Bombus* species with 0.37, 0.38, 3.36 and 15.12 percent of reads mapping to the known viruses or the novel bee viruses found in the study. The poly-A and DSN normalized datasets were unexpectedly highly correlated, given their expected biases (1.000 for *Bombus terrestris*, 0.999 for *Bombus pascuorum* and 0.998 for *Bombus lucorum*) implying that the sequencing results were highly consistent irrespective of the selection method used.

### Previously Described Viruses Present in the Metagenomic Pools

RNAseq reads mapped to three previously described bee viruses. The majority of these reads mapped either to the Acute bee paralysis virus/Kashmir bee virus complex (henceforth ABPV) [29] or to Slow bee paralysis virus (SBPV) [28]. Additionally, in the mixed *Bombus* pool, reads were found mapping to Hubei partiti-like virus 34 (HPLV34) a virus initially detected, though not named, in honeybees by Cornman et al (2012) [30], then subsequently also reported in a sample from Chinese landsnails by Shi et al (2016) [31].

No RNAseq reads were mapped to Deformed wing virus – type A [69], Chronic bee paralysis virus [29], Bee macula-like virus [70], Ganda bee virus [71], Scaldis River bee virus [71], Black queen cell virus [72], Apis rhabdovirus 1 [73], Apis rhabdovirus 2 [73], Apis bunyavirus 1 [73], Apis bunyavirus 2 [73], Apis flavivirus [73], Apis dicistrovirus [73], Apis Nora virus [73] and members of the Lake Sinai virus complex [74]. A small number of small RNA reads did map to these viruses, however, this likely represents cross-mapping, given the lower stringency of 22nt reads. Two of the viral contigs generated by the *de novo* assembly had high similarity to previously described plant viruses; both RNAs of White clover cryptic virus 2 [75] (96% identity), both RNAs of strain of Arabis mosaic virus (MH614320/MH614321) [76] distant to previously sequenced strains (91% identity) and a strain of Red Clover nepovirus A (MH614312) [77] distant to previously sequenced strains (90% identity).

### Putative Novel Viral-like Sequences

We identified 37 putative novel viral contigs, four mapping to DNA viruses (4 densovirus-like contigs) and 33 to RNA viruses (4 Reo group contigs, 2 Toti-Chryso group contigs, 4 Bunya-Arena group contigs, 1 Orthomyxoviridae-like contig, 8 Hepe-Virga group contigs, 12 Picorna-Calici group contigs and 2 Tombus-Noda group contigs). Based on the supposition that a contig represents a separate virus if it maps to a different viral grouping than the other contigs, or if it can be aligned to all other contigs within its assigned viral grouping, this represents 30 new viruses with seven remaining contigs that may represent other genomic regions of these 30 viruses or separate viruses that cannot be confirmed as such. See Table 1 for information on the viruses tested for prevalence using PCR and Supplementary Table 7 for detailed information on all of the identified contigs. The numbers of reads mapping these contigs were variable and are shown in Table 2.

**Table 1.** The names, genome structures and groupings (following Shi et al. (2016) [31]) of the newly discovered viruses for which prevalence was assessed.

| Putative viral contig | Abbreviations | Clade | Genome structure |
| --- | --- | --- | --- |
| Agassiz Rock virus | ARV | Reo | dsRNA |
| Elf Loch virus | ELV | Reo | dsRNA |
| Dumyat virus | DV | Toti-Chryso | dsRNA |
| Sheriffmuir virus | SV | Toti-Chryso | dsRNA |
| Clamshell Cave virus | CCV | Bunya-Arena | - ssRNA |
| Allermuir Hill virus 1 | AHV1 | Hepe-Virga | +ssRNA |
| Allermuir Hill virus 2 | AHV2 | Hepe-Virga | +ssRNA |
| Allermuir Hill virus 3 | AHV3 | Hepe-Virga | +ssRNA |
| Mill Lade virus | MLV | Hepe-Virga | +ssRNA |
| Boghill Burn virus | BBV | Picorna-Calici | +ssRNA |
| Gorebridge virus | GV | Picorna-Calici | +ssRNA |
| Loch Morlich virus | LMV | Picorna-Calici | +ssRNA |
| Mayfield virus 1 | MV1 | Picorna-Calici | +ssRNA |
| Mayfield virus 2 | MV2 | Picorna-Calici | +ssRNA |
| River Liunaeg virus | RLV | Picorna-Calici | +ssRNA |
| Castleton Burn virus | CBV | Tombus-Noda | +ssRNA |

**Table 2.** The RNAseq reads per kilobase per mapped million reads in the *Bombus terrestris, Bombus lucorum, Bombus* pascuorum and mixed *Bombus* pools. Structural zeros are indicated by dashes, zeros in the table indicate below 0.005. Contigs with names in bold meet the criterion of having at least 50 mapping small RNA reads with a sharp peak in the size distribution at 22nt in *Bombus terrestris, Bombus lucorum* and *Bombus pascuorum* providing evidence of replication (see main text).

| Putative viral contig | Accession number | *Bombus terrestris* | *Bombus lucorum* | *Bombus pascuorum* | mixed *Bombus* |
| --- | --- | --- | --- | --- | --- |
| Bombus-associated Densovirus-like Contig 1 | MH614322 | 0.09 | - | - | 12.34 |
| Bombus-associated Densovirus-like Contig 2 | MH614323 | 0.39 | 2.66 | - | 14.23 |
| Agassiz Rock virus | MH614287 | 3.77 | 0.49 | - | - |
| Cnoc Mor virus | MH614297 | 0.97 | - | - | 20.38 |
| Bombus-associated Reoviridae-like Contig 1 | MH614298 | 0.42 | 0.03 | - | 1.63 |
| **Elf Loch virus** | MH614300 | - | 0.00 | 1.00 | 0.05 |
| Dumyat virus | MH614299 | - | - | - | 10.30 |
| Sheriffmuir virus | MH614317 | - | - | - | 2.55 |
| Clamshell Cave virus | MH614294 | 0.07 | - | - | 2.10 |
| Bombus-associated Bunyaviridae-like Contig 1 | MH614295 | 0.70 | - | - | 5.61 |
| Bombus-associated Bunyaviridae-like Contig 2 | MH614296 | - | - | - | 12.13 |
| Bombus-associated Phlebovirus-like Contig 1 | MH614315 | 3.07 | 2.77 | 0.95 | 1.63 |
| Bombus-associated Orthomyxovirus-like Contig 1 | MH614314 | - | - | 0.44 | - |
| **Allermuir Hill virus 1** | MH614288 | 15.27 | 0.64 | 0.02 | 1.50 |
| **Allermuir Hill virus 2** | MH614289 | 0.01 | 0.03 | 12.71 | 0.13 |
| **Allermuir Hill virus 3** | MH614290 | 0.40 | 2.62 | 0.50 | 3.35 |
| **Mill Lade virus** | MH614306 | 0.40 | 0.48 | 0.03 | 7.28 |
| Bombus-associated Virga-like Contig 1 | MH614308 | - | 0.65 | - | - |
| Bombus-associated Virga-like Contig 2 | MH614309 | - | 0.33 | - | - |
| Bombus-associated Virga-like Contig 3 | MH614318 | 16.03 | 4.83 | 0.52 | 90.37 |
| Bombus-associated Virga-like Contig 4 | MH614319 | 1.66 | 0.11 | - | 0.54 |
| Black Hill virus | MH614291 | - | - | - | 2.96 |
| Boghill Burn virus | MH614292 | 0.00 | 10.92 | 0.00 | 0.11 |
| Gorebridge virus | MH614301 | 2.97 | 0.02 | - | 0.15 |
| Bombus-associated Picornavirus-like Contig 1 | MH614302 | 6.27 | 0.02 | - | 0.29 |
| Loch Morlich virus | MH614303 | 0.00 | - | - | 7.83 |
| **Mayfield virus 1** | MH614304 | 391.31 | 232.93 | 0.59 | 7.67 |
| **Mayfield virus 2** | MH614305 | 4.87 | 0.79 | 336.97 | 558.82 |
| Bombus-associated Nepovirus-like Contig 1 | MH614310 | 1.72 | 1.02 | 0.09 | 1.70 |
| Bombus-associated Nepovirus-like Contig 2 | MH614311 | 0.33 | 0.39 | 0.11 | 0.21 |
| Bombus-associated Picornavirus-like Contig 2 | MH614316 | 0.11 | 0.29 | 1.02 | 0.04 |
| River Liunaeg virus | MH614307 | 0.24 | 0.47 | 0.01 | 9.58 |
| **Castleton Burn virus** | MH614293 | 2.39 | 12.30 | 8.10 | 122.99 |
| Bombus-associated Nodavirus-like Contig 1 | MH614313 | - | - | - | 1.50 |

### siRNA-based Evidence for Infection

RNA interference is an important component of antiviral defence in arthropods [78]. As part of this defence mechanism, homologs of *Drosophila* Dicer-2 cleave dsRNA, usually in the form of replication intermediates, giving rise to a characteristically narrow and sharply peaked distribution of virus-derived small RNAs. Thus the presence of such small RNAs from both strands of an ssRNA virus provide compelling evidence that the virus was replicating. In bumblebees the characteristic Dicer-mediated viral siRNAs peak sharply at 22nt [73], and the viruses that displayed at least 50 characteristic viral siRNAs are marked in Table 2. The distribution of the mapped small RNA reads is shown in Fig 3 for all viruses where the siRNAs are described in the main text, with full data in Supp Fig 1.

**Fig 3.**
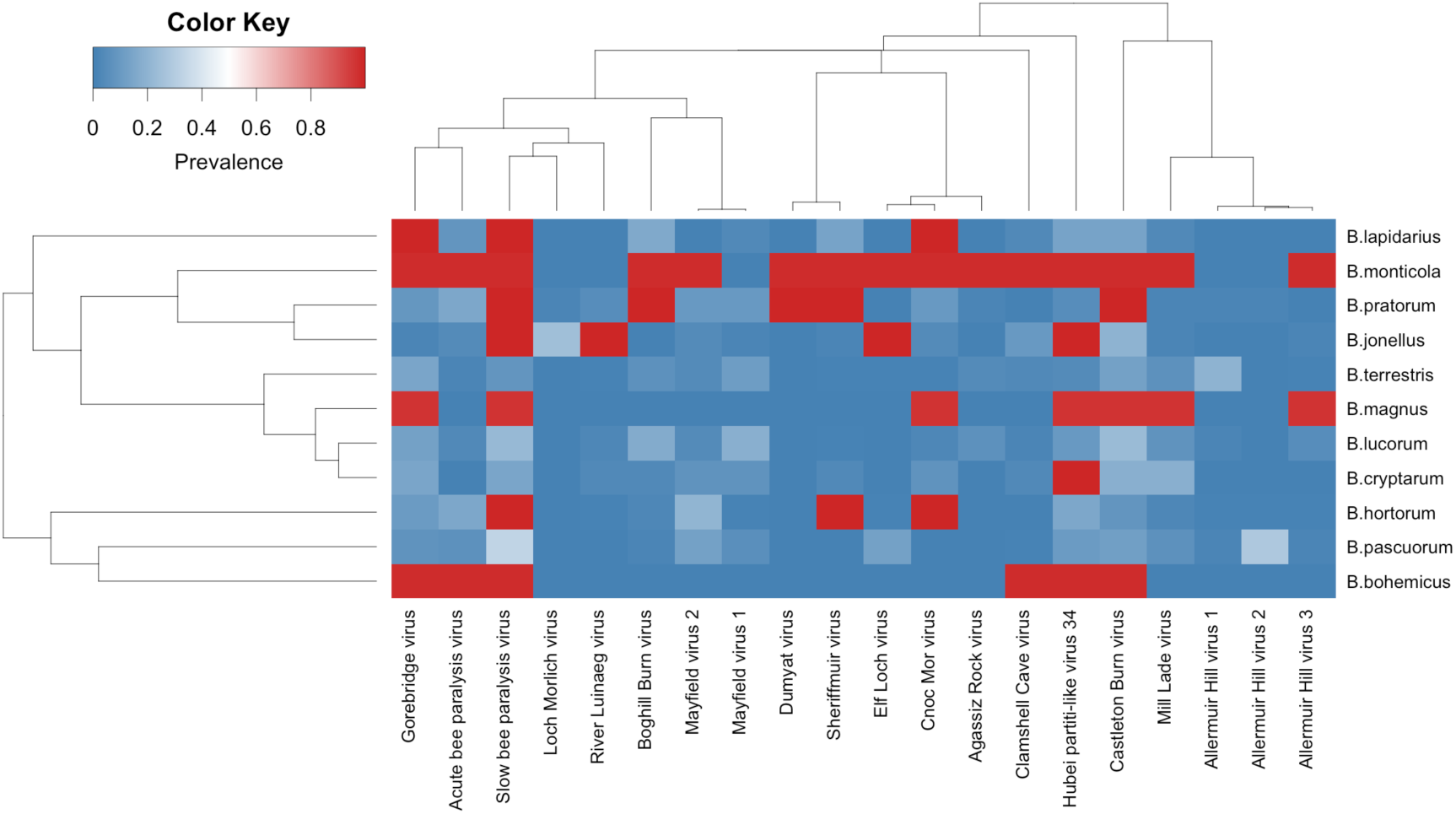
A heatmap of maximum likelihood estimates for prevalence. Hosts and viruses are ordered by phylogenetic relatedness, the trees represent the maximum clade credibility topology. Squares in red with maximum likelihood estimates of the prevalence of 1 correspond to cases where all pools were positive. The maximum likelihood estimate is likely extremely upwardly biased in this case.

In the three bumblebee species with siRNA data, a sequence similar to a phlebovirus glycoprotein (AEL29653.1) displayed >50 siRNA reads. However, the size spectra of these reads is centered on 24nt with a strong bias for a 5’ terminal uracil, with the antisense mapping orientation being more prevalent. This 5’ U-bias is consistent with insect piRNAs [79], and the predominant antisense orientation is consistent with the piRNA mapping pattern to endogenous viral elements (EVE) in mosquitoes [80]. However, the size of piRNAs in bumblebees is generally larger than this [81]. This sequence is therefore potentially an EVE that has either been gained multiple times or has been maintained in the bumblebee genome since at least the *B. pascuorum*-*B. terrestris*/*B. lucorum* split.

It is notable that the size distribution of viral siRNAs is less sharply peaked in Mayfield virus 1, Mayfield virus 2 and Slow bee paralysis virus (excepting Mayfield virus 1 in *B. lucorum*, which is sharply peaked), with broad ‘shoulders’. This is reminiscent of the pattern seen for Drosophila C virus and Drosophila Nora virus in wild-collected D. *melanogaster* [61], both of which contain a viral suppressor of RNAi [82,83].

*B. pascuorum* also had siRNA reads mapping to a sequence with 90% identity to Red clover nepovirus A (MG253829.1) [77]. However, the read length spectra were sharply peaked at 21nt, rather than the 22nt of bumblebee viRNAs. This is consistent with siRNA’s produced from DLC4, the key antiviral dicer in *Arabidopsis thaliana* [84] implying acquisition of the small RNAs through nectar or pollen contamination.

### Prevalence

Species level prevalences differed dramatically among the different viruses (Fig 3). Prevalences were generally low to intermediate, with modal viral prevalences for most host-virus combinations being below 15%. Slow bee paralysis virus was by far the most common virus in the sample, with estimated prevalences of greater than 25% in multiple species. Our ability to estimate the prevalence of common viruses is limited by the pooling, leading us to only be able to assign lower bounds to prevalences in these cases, but in 7 of 11 species, all pools were positive for SBPV. Acute bee paralysis virus, Hubei partiti-like virus 34, Castleton Burn virus, Gorebridge virus, Mayfield virus 1 and Mayfield virus 2 all reached 15-25% prevalences in multiple species. Several viruses showed strong signals of species specificity, having very low to zero prevalences in multiple host species but high prevalences in others. Examples of this pattern include Allermuir Hill virus 1 in *B. terrestris*, Allermuir Hill virus 2 in *B. pascuorum*, Allermuir Hill virus 3 in *B. magnus* and *B. monticola*, as well as Loch Morlich virus and River Luinaeg virus in *B. jonellus*.

### Host-Pathogen Co-phylogenetic Models

All models that included a virus phylogeny term gave qualitatively similar results (Fig 4, Table 3). This suggests that the results are robust to both phylogenetic uncertainty and the assumption of a common ancestor of all RNA viruses. For this reason, for the rest of this section, estimates will be given from the model containing the estimated phylogeny with all the RNA viruses included. All estimates represent the percentage of the total variance in the model (the sum of all estimated variance components adjusted for the variance of the link function by the addition of π^2^/3) explained by a term. In all cases, the presented point estimate is the posterior mean, and 90% shortest posterior density intervals [66] are presented following in square brackets. Shortest posterior density intervals are a variant of highest posterior density intervals and describe the shortest possible interval containing (in this case) 90% of the probability density for the parameter. We present 90% intervals rather than the standard 95% intervals, as 95% intervals calculated from simulation draws are less computationally stable [67]. In all cases but the virus phylogenetic effect, the posterior estimates for the proportion of variance explained by each effect differed strongly from their induced priors (Supp Fig 2).

**Fig 4.**
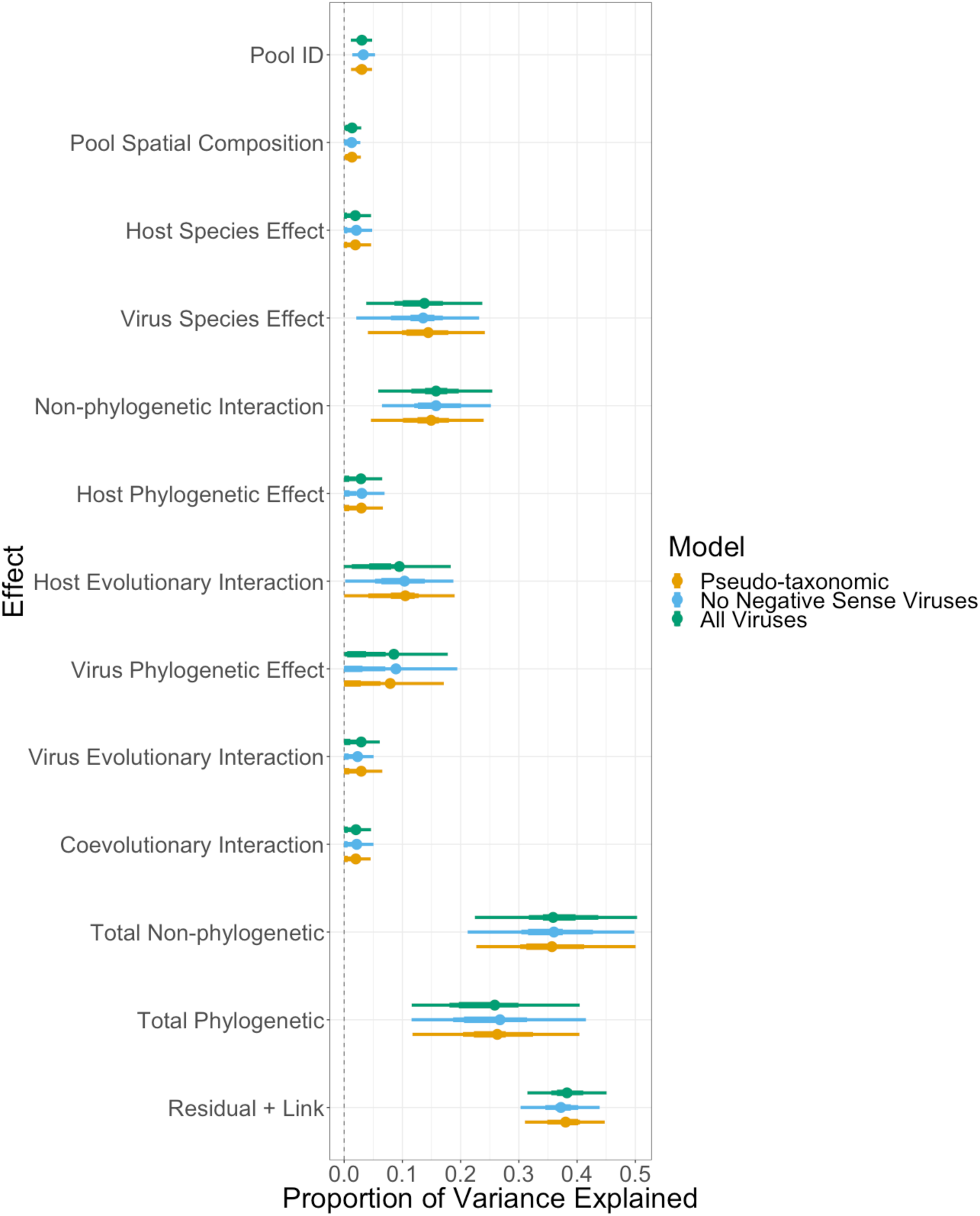
Comparison of estimated proportion of variance in prevalence explained by different parameters between models. As each factor explains a proportion of the attributed variance in the model, total variance over all factors must sum to 1. For each parameter, the circle represents the modal estimate, the thick bars represent the 50% shortest posterior density interval and the thin bars represent the 90% shortest posterior density interval. “Pool ID” is the proportion of the total variation in prevalence explained by pools within species differing in the degree to which they were infected by viruses. “Spatial Composition” is the proportion of the total variation in prevalence explained by the combination of locations from which the bees in the pool originate. “Host Species Effect” is the proportion of variation in prevalence explained by hosts having different average viral prevalences. “Virus Species Effect” is the proportion of variation in prevalence explained by viruses differing in their average prevalences. “Non-phylogenetic Interaction” is the proportion of variation in prevalence explained by host-virus combinations differing in the their average prevalences beyond that which would be expected by their host and virus species effects alone. “Host Phylogenetic Effect” is the proportion of variation in prevalence explained by hosts having average viral prevalences correlated across the host phylogeny. “Virus Phylogenetic Effect” is the proportion of variation in prevalence explained by viruses having average prevalences correlated across the viral phylogeny. “Host Evolutionary Interaction” is the proportion of variation in prevalence explained by related hosts having correlated viral assemblages. “Virus Evolutionary Interaction” is the proportion of the variation explained by related viruses having correlated host assemblages. “Coevolutionary Interaction” is the proportion of the variation explained by related hosts having similar prevalences of related viruses. “Total Non-phylogenetic” is the proportion of the variation that can be explained by terms not involving the host and virus phylogeny and excluding the residual (“Host Species Effect”, “Virus Species Effect”, “Pool ID”, “Spatial Composition Effect”, “Non-phylogenetic Interaction”). “Total phylogenetic” is the proportion of the variation that can be explained by terms involving a host or virus phylogeny (“Host Phylogenetic Effect”, “Virus Phylogenetic Effect”, “Host Evolutionary Interaction”, “Virus Evolutionary Interaction”, “Coevolutionary Interaction”). “Residual + Link” is the proportion of the total variance that is explained by the residual variance and variance of the logistic distribution (π^2^/3).

**Table 3.**
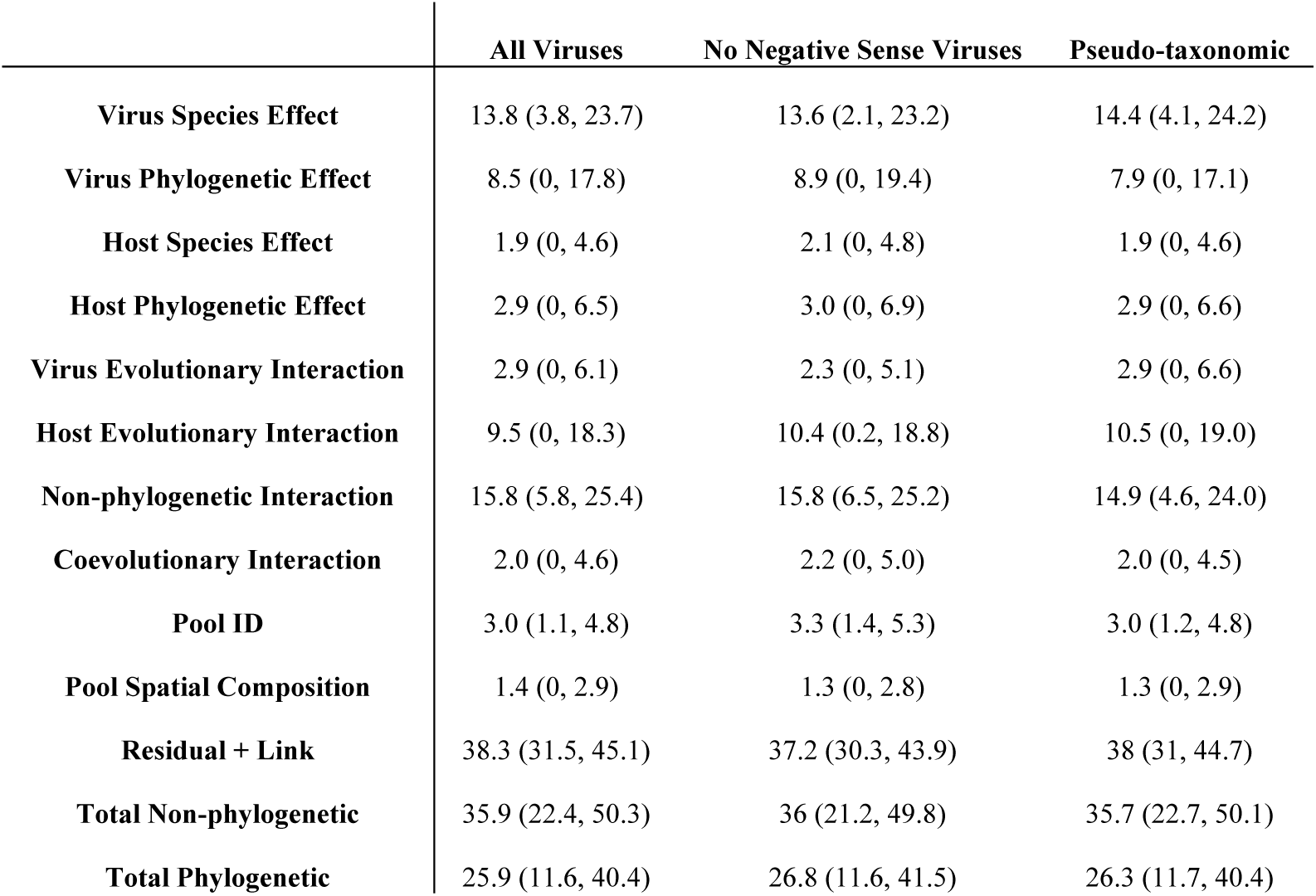
Mean estimates for the intra-class correlations of each variance component. The point estimate is the posterior mean, the numbers in brackets represent the 90% shortest posterior density interval.

### Summary of Model Results

We find evidence that which viral species infects a host, the specific interaction between individual hosts and individual viruses and related hosts having similar prevalences with the same sets of viruses all explain variation in infection prevalence.

### Total Evolutionarily-associated Variation

In the models containing the virus phylogeny, approximately a quarter (25.9% [11.6-40.4]) of the total variation in prevalence was explained by terms accounting for the evolutionary histories of hosts and viruses (host phylogenetic effect, virus phylogenetic effect, host evolutionary interaction effect, virus evolutionary interaction effect and coevolutionary interaction).

### Host and Virus Level Effects

The host and virus phylogenetic effects measure the extent to which related hosts have similar average prevalences of virus infection and related viruses have similar average prevalences across hosts. The host and virus non-phylogenetic terms measure the extent to which hosts and viruses differ in their average infection levels in manner not consistent with evolution by Brownian motion along a phylogeny. The host species and phylogenetic effects explained a small proportion of the total variance in infection probability (species: 1.9%, [0.0-4.6]; phylogenetic: 2.9%, [0.0-6.5]). The shape of the posterior distributions for the two parameters visualised in Fig 4, makes it clear that the most credible values for both of these parameters are 0. While it is unlikely that there is no variation in average prevalence between host species, it is clear that the amount of prevalence explained by hosts differing in their infection levels averaged across viruses is small relative to the other effects.

The virus species and phylogenetic effects explained a larger proportion of the total variance in infection probability, with non-phylogenetic variation dominating, but were imprecisely estimated (species: 13.8%, [3.8-23.7]; phylogenetic: 8.5%, [0.0-17.8]). The posterior density for the phylogenetic effect is concentrated at 0. So, while it is clear there is a virus species effect, the data does not appear informative for the presence or absence of a viral phylogenetic effect in this system. The posterior draws for the viral species and phylogenetic effect were negatively correlated within iterations, indicating that the model had difficulty partitioning the two. This partial non-identifiablility explains the broad posteriors on both.

### Interaction Effects

A host evolutionary interaction effect measures the extent to which more closely related hosts have more similar prevalences with the same sets of viruses, and a virus evolutionary interaction effect measures the extent to which more closely related viruses infect the same sets of hosts to more similar degrees. A coevolutionary interaction term measures the extent to which more related hosts are infected to similar degrees with viruses that are themselves related. The non-phylogenetic interaction term measures the extent to which there is variance in the mean prevalence of specific host-virus pairings, that are is not consistent with the other interaction effects (see Box 1).

There was little evidence for a virus evolutionary interaction effect or coevolutionary interaction having a large effect on the observed prevalences (virus: 2.9% [0.0-6.1]; coevolutionary: 2.0% [0.0-4.6]). In both cases, the marginal posterior distributions peaked at 0.

There was evidence for a host evolutionary interaction explaining some of the total variance in prevalence (9.5% [0.0-18.3]). This is the only parameter in the model for which the estimated size of the effect depended strongly on the specific treatment of the virus phylogeny (see Fig 4). Whether the lower 90% bound of the credible interval for the variation explained rounded to 0.0 or 0.1 depended on the phylogenetic matrix (or set of phylogenetic matrices) inputted. The marginal posterior distribution of the parameter was concentrated at lower values when the estimated phylogeny including the negative-sense RNA viruses was used, and at higher values in the other two cases. However, irrespective of the choice of virus phylogeny, the mode of the distribution and majority of the density was distant from zero, implying that the effect is likely to be biologically relevant. As the virus phylogenies themselves are not actually directly involved in this term, this must be due to the partitioning of variance across other terms being cryptically different depending on the assumptions about the virus phylogeny.

There was also a clear non-phylogenetic interaction (15.8% [5.8-25.4]), implying that much of the variation in prevalence is due to the idiosyncrasies of individual host-virus combinations.

As with the virus species effect and the virus phylogenetic effect, the MCMC draws for the proportion of variance explained by the host evolutionary interaction and non-phylogenetic interaction were negatively correlated within an iteration, implying that separating these parameters was proving difficult. While this led to a diffuse posterior with wide credible intervals for both, they remain individually interpretable, and both effects appear present simultaneously.

### Tongue Length-Viral Community Correlation

Given that the co-phylogenetic model found that related hosts share viral communities, one potential mechanism for this is phylogenetically-biased exposure, driven by phylogenetically correlated floral preferences. If bumblebee species with similar flower preferences had similar viral communities, it would be expected that there would be a positive correlation between tongue length similarity (as this is an important factor in floral preference) and viral community similarity between pairs of species. The point estimate of the correlation between the two distances was small and negative (−0.06), but the 95% confidence intervals for that point estimate overlapped zero (−0.13, 0.00), so given the uncertainty in the data, a correlation of zero cannot be rejected. Nonetheless, given this data, a strong positive relationship between tongue length and viral community similarity seems unlikely, a result inconsistent with phylogenetically-biased exposure driven by tongue length-mediated floral choice.

## Discussion

Using wild bumblebee species that share transmission opportunities, we have shown that variation in the prevalence of infection in the wild is explained by related hosts being infected with the same viruses to similar degrees, viruses differing in their average prevalence and individual virus-host pairings having greater or lesser prevalence than would be expected by the host and virus species effects alone.

### Virus Discovery

There is now an extensive diversity of viruses known in bees, with most new studies finding novel viruses [30,71,73,74,85-87]. We have found 37 novel putative viral contigs in the transcriptomes of wild-caught bumblebees from across Scotland, suggesting that virus discovery in this taxonomic group is far from saturation. As with any metagenomic study, it is hard to be confident that the virus-like contigs represent real infections of the sampled host, rather than surface or gut contaminants. However, the presence of 22nt virus siRNAs, generated from double-stranded viruses by Dicer as part of an antiviral response in the host, provides compelling evidence that at least 10 of these contigs (Densovirus 2 and 3, Elf Loch virus, Allermuir Hill virus 1, 2 and 3, Mill Lade virus, Mayfield virus 1 and 2, and Castleton Burn virus) represent active viral infections in bumblebees.

Mites and nematodes both parasitise bumblebees and therefore could potentially be an alternative source of the small RNAs. Mite viRNAs are reported to be centered at 24nt [73], and could therefore not produce the small RNA patterns observed. Nematode viRNAs are centered at 22nt, like bumblebee viRNAs, [88] and thus could potentially produce this pattern. While, outside of queens infected with *Sphaerularia bombi*, nematode infection of wild bumblebees appears to be very rare [89], nematodes cannot be categorically ruled out as a source of the observed small RNAs. One contig’s (MH614312) mapped small RNAs were centered at 21nt, and the closest known virus was a nepovirus of plants. As DCL4, a major plant Dicer, produces viRNAs of this size [90], this is consistent with that particular virus being a plant virus, which was transferred in collected nectar or pollen.

### Phylogenetic Effects

We found no evidence for a large host phylogenetic effect, where related hosts have correlated average viral prevalences. To the best of our knowledge no studies have previously applied these methods to viruses sampled from wild animals. However, other traits relating to viral disease in a series of studies in *Drosophila* species under experimental conditions have consistently detected host phylogenetic effects in factors that would be expected to be correlated with prevalence in the wild, such as infection probability [24], virulence and viral load [91] and viral load alone [92]. However, two of these studies focused on a single isolate of Drosophila C virus. Therefore, the variation that they attribute to a phylogenetic effect may be partitioned into the host evolutionary interaction in our study, as a host evolutionary interaction is equivalent to separate inconsistent host phylogenetic effects for each virus. Our data were not particularly informative for the presence or absence of a virus phylogenetic effect, with the posterior being very diffuse with a majority of the density near zero. This appears partly due to difficulties partitioning the variation between the virus species effect and virus phylogenetic effect. Irrespective of the cause, we can make no strong statements about whether related viruses exhibit similar prevalences across hosts from this dataset.

### Host Species Effect

There was little evidence for an important effect of host species independent of the phylogeny, implying that hosts do not strongly vary in the average degree to which they are infected with viruses. The previous studies using these methods have universally found host species effects [25,93]. However, as both of these studies have used mammal-eukaryotic parasite datasets, the degree of relevance for them as a comparison in unclear. Experimental evidence from virus studies across drosophilid flies have found weak to zero host species effects on the titre of sigma viruses [24] and Drosophila C virus virulence [91] but considerably larger host species effects on Drosophila C virus load [91]. This between-study variation potentially indicates a reason we did not detect a host species effect. With a single pathogen, the average and particular degrees of variation in infection between hosts are identical. As soon as multiple pathogens are involved, they diverge, such that it is possible for there to be no variation in the average prevalence between hosts, but still considerable variation in the prevalence of particular viruses between hosts, which is consistent with the presence of a host species effect in correlates of prevalence in some viruses but not others, as noted above.

### Virus Species Effect

A clear effect of virus species independent of the virus phylogeny was detected in the dataset, despite the uncertainty added by the difficulty partitioning the virus species and phylogenetic effects. Therefore, viruses differed in their prevalences averaged across hosts. This is an intuitive result as viruses differ in host range [94], virulence [95,96] and infectious period [96] at both the species and the strain level. Variation in host range changes the size of the host pool available for infection, and variation in virulence and infectious period both change the length of time any infected host is available for sampling. All these factors would be expected to drive consistent differences in long-run prevalences between viruses. Additionally, as our sites were only sampled once, short-term effects will also drive between virus variation. Any virus that was experiencing an epizootic at the time of sampling will be overrepresented relative to its long-run prevalence, further increasing the between virus variation.

### Host Evolutionary Interaction Effect

A host assemblage effect was found, where phylogenetically related hosts share viral assemblages, showing that more closely related hosts are more similar in virus prevalence for groups of viruses. The statistical machinery required for estimating this effect is quite new and, in the disease ecology field, has predominantly been applied to mammal-parasite and plant-parasite systems where there are good datasets already. Nonetheless, host evolutionary interactions have always been found when searched for using these methods [25,93] and analogous effects are commonly found using different methods [97–99]. In a system where these viruses were host limited, this pattern could be explained by preferential host shifting, where parasites more frequently gain the ability to infect hosts closely related to ones that they are already capable of infecting. Preferential host shifting is known to be a general phenomenon, and has been observed in macroparasites, viruses and protozoans (see Longdon et al (2014) [27] and the references within).

While some of the viruses in this study were not detected in a subset of host species, most of the viruses found here appear to genuinely be multihost viruses, with the majority being detected in over half the sampled species. Given this, a combination of biased cross-species transmission and preferential host shifting appears a better explanation in this system. Biased cross-species transmission occurs when transmission occurs more frequently between some species that a pathogen is already capable of infecting than others. This biased cross-species transmission could be driven by two non-exclusive mechanisms: phylogenetically-biased transmission probabilities and phylogenetically-biased exposure.

Phylogenetically-biased transmission probabilities occur when cross-species transmission is more frequent among close relatives, due to the probability of infection after contact with the virus being similar between related species. Related hosts present correlated environments from the perspective of the virus at the molecular and anatomical level, therefore adaptation to one should provide corresponding fitness increases on the other. Experimental results have shown that correlated mutations occur on viral entry to related hosts [100], implying that this cross-adaptation does occur. However, this is probabilistic, and the mutations fixed on entry can differ between replicate entries [100,101]. Therefore, if there is antagonistic pleiotropy between mutations that are adaptive in two different groups of hosts and cross-adaptation predicts the probability of successful infection on contact, then a phylogenetically-biased transmission network will result.

Phylogenetically-biased exposure represents an evolutionarily-driven ecological phenomenon that biases cross-species transmission rates, mediated by niche overlap. Contaminated flowers are likely to be an important source of intra- and inter-specific pathogen transmission in bumblebees and pollinators more generally [102–104]. The flower visitation network has been shown to be associated with the partitioning of genetic diversity of *Crithidia bombi* between bumblebee hosts [9], and the network itself is highly structured, though temporally variable [105]. Different bumblebee species show tongue length differences, which are phylogenetically associated [15], and the differences in tongue length correlate with differential flower usage between bumblebee species [12,106]. If infection occurs at contaminated flowers, the structuring of the flower usage network could cause different flowers to build up different surface viral communities. This could drive consistent phylogenetically-correlated differences in viral infection rates through differential exposure.

Post-hoc testing did not find a positive relationship between tongue length dissimilarity (a rough proxy for species-level flower choice dissimilarity) and viral community composition dissimilarity, which provides some evidence against phylogenetically-biased exposure as the causative mechanism. However, the study design in this case is not optimal for disentangling biased transmission probabilities and biased exposure, as species were sampled from different locations at a single timepoint and prevalence of the viruses varied spatially. Given this, drawing strong conclusions as to the relative impact of the two mechanisms outlined above based on this data would be premature. Similarly, the subgenus *Psithyrus* contains socially parasitic species that are coevolved to parasitise particular social bumblebee species, which could also lead to phylogenetically-correlated differences in viral infection rates. We were unable to test whether socially parasitic cuckoo bumblebee species have similar viral communities to their hosts, as our study included only a single parasitic bumblebee species, *B. bohemicus*, but the possibility of brood parasitism being an important driver of between-colony disease transmission is worth further study.

### Virus Evolutionary Interaction Effect and Coevolutionary Interaction

We found no evidence of a large virus evolutionary interaction or coevolutionary interaction. This is largely unsurprising as it would appear implausible that the host assemblages have been conserved over evolutionary time, as the deep splits in the viral families predate the most recent common ancestor of bumblebees by many millions of years [107].

### Non-phylogenetic Interaction

A non-phylogenetic interaction was detected. This interaction represents variation in prevalence caused by specific host-virus pairings having prevalences beyond that which would be expected by the simple addition of the individual host and virus means. A non-phylogenetic interaction could be caused by a large range of factors, some biological and some due to the specifics of the model, many of which would be likely to be acting simultaneously to generate this signal. One possibility is coevolution between the host and virus that occurred after both diverged from their common ancestor with the closest related species in the study. Another is the complete absence of coevolution, where spillover from a primary or group of primary hosts causes either a constant very low prevalence of dead-end infections, which are none-the-less detectable by PCR. Related to this is a statistical issue involving cases where not every species in the study is within a virus’ host range, and the species that are within the host range are not closely related. In this case, the variation is not absorbed by the host evolutionary interaction and almost no host has a prevalence close to the mean across hosts, as in many species the prevalence is zero, which causes the mean to be considerably lower than the average prevalence in the species the virus does infect. This effect would be magnified if the sampling occurs during an epizootic. More broadly, anything that changes the epidemiological parameters of a virus in a specific host will lead to a non-phylogenetic interaction. Considering the variation in the natural history of viruses and the lesser, but still significant, variation in the natural history of bumblebees, a large non-phylogenetic interaction is to be expected.

## Conclusion

While it is clear that viruses are abundant in pollinators, the factors that determine the distribution of pollinator viruses have remained uncertain, outside of a few well-studied cases [108,109]. With the novel viruses discovered in this study, we have investigated predictors of these virus/host associations and found that both the host evolutionary history and the identity of the virus contributes to this distribution. This supports both theory and prior empirical evidence that related species are more at risk of infection from each other’s diseases than the diseases of distantly related species. However, the importance of the viral identity and unique interactions between host-virus pairs suggests that the introduction of a novel virus into a community is likely to have unpredictable effects even when no close relatives of currently known hosts are present. This highlights the risk posed by disease spillover for the conservation not only of wild pollinator communities, but also to communities consisting of related animal or plant species in general.

## Supporting information

All supplementary materials

## Data Availability

The data and code for running the analyses is available on github under a GPLv3 licence, as code uses code taken from other GPLv3 licenced works (https://github.com/dpascall/bumblebee-virus-cophylo).

## Acknowledgements

We thank Jarrod Hadfield for extensive statistical advice, modifications to MCMCglmm and for helpful comments on this manuscript, Dave Goulson for assistance identifying some specimens and to Ben Longdon and Bethany Clark for further comments. Rowan Doff assisted in the lab with RFLP analyses and Claire Webster with DNAse treatments. Claire Webster, Jarrod Hadfield and Florian Bayer helped with fieldwork.

## Funding

This work was funded by a BBSRC SWBIO DTP PhD stipend to DP, a Royal Society Dorothy Hodgkin Fellowship to LW, Wellcome Trust Research Career Development Fellowship WT085064 to DJO.

Author Contributions
DJP – Conceptualization, Data Curation, Formal 24 Analysis, Investigation, Methodology, Project Administration, Software, Visualization, Writing – Original Draft Preparation, Writing – Review & Editing
MCT – Investigation, Resources, Writing – Review & Editing
DJO – Conceptualization, Data Curation, Formal Analysis, Funding Acquisition, Methodology, Project Administration, Software, Visualisation, Writing – Review & Editing
LW – Conceptualization, Funding Acquisition, Investigation, Methodology, Project Administration, Resources, Supervision, Writing – Review & Editing

