## Supplementary material for "Host evolutionary history predicts virus prevalence across bumblebee species": All supplementary materials

Supplementary Table 1 The longitudes and latitudes of the sites sampled

| **Location** | **Latitude** | **Longitude** |
| --- | --- | --- |
| Dalwhinnie | 56.935 | -4.245 |
| Edinburgh | 55.921 | -3.179 |
| Glenmore | 57.166 | -3.696 |
| Gorebridge | 55.826 | -3.046 |
| Iona | 56.328 | -6.406 |
| Ochils | 56.158 | -3.88 |
| Pentlands | 55.796 | -3.404 |
| Staffa | 56.436 | -6.341 |
| Stirling | 56.149 | -3.916 |

Supplementary Table 2 The number of bees in each pool and their species

| **Pool ID** | **Species** | **Sample Size** | **Location** |
| --- | --- | --- | --- |
| BO | *Bombus bohemicus* | 10 | Gorebridge, the Ochils, the Pentlands and Glenmore |
| C1 | *Bombus cryptarum* | 10 | Ochills |
| C2 | *Bombus cryptarum* | 10 | Ochills |
| C3 | *Bombus cryptarum* | 5 | Iona, the Ochils and the Pentlands |
| C4 | *Bombus cryptarum* | 8 | Gorebridge and Edinburgh |
| DIV | Mixed | 7 | Gorebridge, Staffa, the Ochils and the Pentlands |
| H1 | *Bombus hortorum* | 10 | Gorebridge |
| H2 | *Bombus hortorum* | 10 | Gorebridge |
| H3 | *Bombus hortorum* | 10 | Gorebridge |
| H4 | *Bombus hortorum* | 10 | Gorebridge |
| H5 | *Bombus hortorum* | 10 | Gorebridge |
| H6 | *Bombus hortorum* | 6 | Gorebridge |
| H7 | *Bombus hortorum* | 7 | Gorebridge |
| H8 | *Bombus hortorum* | 7 | Stirling, Glenmore and Dalwhinnie |
| H9 | *Bombus hortorum* | 11 | Iona |
| J1 | *Bombus jonellus* | 10 | Stafa |
| J2 | *Bombus jonellus* | 10 | Stafa |
| J3 | *Bombus jonellus* | 5 | Iona and Staffa |
| J4 | *Bombus jonellus* | 10 | Glenmore |
| J5 | *Bombus jonellus* | 10 | Glenmore |
| J6 | *Bombus jonellus* | 10 | Glenmore |
| J7 | *Bombus jonellus* | 5 | Dalwhinnie and Glenmore |
| L1 | *Bombus lucorum* | 10 | Stirling and the Ochils |
| L2 | *Bombus lucorum* | 10 | Stirling and the Ochils |
| L3 | *Bombus lucorum* | 10 | Gorebridge |
| L4 | *Bombus lucorum* | 10 | Gorebridge |
| L5 | *Bombus lucorum* | 10 | Gorebridge |
| L6 | *Bombus lucorum* | 10 | Gorebridge |
| L7 | *Bombus lucorum* | 10 | Gorebridge |
| L8 | *Bombus lucorum* | 10 | Gorebridge |
| L9 | *Bombus lucorum* | 10 | Gorebridge |
| L10 | *Bombus lucorum* | 10 | Gorebridge |
| L11 | *Bombus lucorum* | 8 | Gorebridge |
| L12 | *Bombus lucorum* | 10 | Edinburgh |
| L13 | *Bombus lucorum* | 10 | Edinburgh |
| L14 | *Bombus lucorum* | 10 | Edinburgh |
| L15 | *Bombus lucorum* | 10 | Stirling, the Ochils and the Pentlands |
| L16 | *Bombus lucorum* | 10 | Iona, the Ochils and the Pentlands |
| L17 | *Bombus lucorum* | 2 | The Ochils |
| L18 | *Bombus lucorum* | 10 | Edinburgh |
| L19 | *Bombus lucorum* | 10 | Edinburgh |
| L20 | *Bombus lucorum* | 2 | Edinburgh |
| LA1 | *Bombus lapidarius* | 11 | Gorebridge |
| LA2 | *Bombus lapidarius* | 8 | Gorebridge |
| LA3 | *Bombus lapidarius* | 8 | Edinburgh |
| LA4 | *Bombus lapidarius* | 7 | Stirling and the Ochils |
| M | *Bombus monticola* | 10 | Glenmore, the Ochils and the Pentlands |
| MA | *Bombus magnus* | 11 | Iona, the Ochils and Glenmore |
| P1 | *Bombus pascuorum* | 10 | Stirling |
| P2 | *Bombus pascuorum* | 10 | Ochills |
| P3 | *Bombus pascuorum* | 9 | Iona, Staffa and the Pentlands |
| P4 | *Bombus pascuorum* | 10 | Ochills |
| P5 | *Bombus pascuorum* | 10 | Ochills |
| P6 | *Bombus pascuorum* | 10 | Ochills |
| P7 | *Bombus pascuorum* | 8 | Stirling |
| P8 | *Bombus pascuorum* | 10 | Ochills |
| P9 | *Bombus pascuorum* | 5 | Ochills |
| P10 | *Bombus pascuorum* | 10 | Edinburgh |
| P11 | *Bombus pascuorum* | 10 | Edinburgh |
| P12 | *Bombus pascuorum* | 10 | Edinburgh |
| P13 | *Bombus pascuorum* | 10 | Edinburgh |
| P14 | *Bombus pascuorum* | 10 | Edinburgh |
| P15 | *Bombus pascuorum* | 5 | Edinburgh |
| P16 | *Bombus pascuorum* | 10 | Gorebridge |
| P17 | *Bombus pascuorum* | 10 | Gorebridge |
| P18 | *Bombus pascuorum* | 10 | Gorebridge |
| P19 | *Bombus pascuorum* | 10 | Gorebridge |
| P20 | *Bombus pascuorum* | 10 | Gorebridge |
| P21 | *Bombus pascuorum* | 10 | Gorebridge |
| P22 | *Bombus pascuorum* | 10 | Gorebridge |
| P23 | *Bombus pascuorum* | 5 | Gorebridge |
| PR1 | *Bombus pratorum* | 10 | Edinburgh |
| PR2 | *Bombus pratorum* | 10 | Edinburgh |
| PR3 | *Bombus pratorum* | 9 | Edinburgh |
| PR4 | *Bombus pratorum* | 10 | Gorebridge |
| PR5 | *Bombus pratorum* | 8 | Gorebridge, Stirling and the Ochils |
| T1 | *Bombus terrestris* | 10 | Stirling and the Ochils |
| T2 | *Bombus terrestris* | 10 | Gorebridge |
| T3 | *Bombus terrestris* | 10 | Gorebridge |
| T4 | *Bombus terrestris* | 10 | Gorebridge |
| T5 | *Bombus terrestris* | 10 | Gorebridge |
| T6 | *Bombus terrestris* | 10 | Gorebridge |
| T7 | *Bombus terrestris* | 10 | Gorebridge |
| T8 | *Bombus terrestris* | 10 | Gorebridge and Edinburgh |
| T9 | *Bombus terrestris* | 10 | Edinburgh |
| T10 | *Bombus terrestris* | 10 | Edinburgh |
| T11 | *Bombus terrestris* | 10 | Edinburgh |
| T12 | *Bombus terrestris* | 10 | Edinburgh |
| T13 | *Bombus terrestris* | 10 | Edinburgh |
| T14 | *Bombus terrestris* | 10 | Stirling and the Ochils |
| T15 | *Bombus terrestris* | 10 | Stirling and the Ochils |
| T16 | *Bombus terrestris* | 5 | Stirling and the Ochils |
| T17 | *Bombus terrestris* | 10 | Gorebridge |
| T18 | *Bombus terrestris* | 10 | Gorebridge |
| T19 | *Bombus terrestris* | 8 | Gorebridge |
| T20 | *Bombus terrestris* | 10 | Edinburgh |
| T21 | *Bombus terrestris* | 10 | Edinburgh |
| T22 | *Bombus terrestris* | 6 | Edinburgh |
| T23 | *Bombus terrestris* | 10 | Gorebridge |
| T24 | *Bombus terrestris* | 10 | Gorebridge |
| T25 | *Bombus terrestris* | 10 | Gorebridge |
| T26 | *Bombus terrestris* | 10 | Gorebridge |

Supplementary Table 3 The PCR conditions for all reactions in this paper. Codes in the conditions box refer to below

PCR 62: 95**°**C 5:00, (95**°**C 0:15, 62**°**C 0:30, 72**°**C 0:45)x40, 72**°**C 7:00, 4**°**C

PCR 60: 95**°**C 5:00, (95**°**C 0:15, 60**°**C 0:30, 72**°**C 0:45)x40, 72**°**C 7:00, 4**°**C

T62: 95**°**C 5:00, (95**°**C 0:15, 62**°**C [minus 1**°**C per step] 0:30, 72**°**C 0:45)x10, (95**°**C 0:15, 52**°**C 0:30, 72**°**C 0:45)x30, 72**°**C 7:00, 4**°** C

| **Primer** | **Virus** | **Sequence** | **Conditions** | **Product Size** | **Reference** |
| --- | --- | --- | --- | --- | --- |
| MV1Forward | Mayfield virus 1 | TATCCGCCGGCGTAATCTTC | PCR 60 | 481 |  |
| MV1Reverse | Mayfield virus 1 | GGATCTGATCCGTAGCGTGG |  |  |  |
| MV2Forward | Mayfield virus 2 | CGGCTGCGTTGCGTAGTATA | T62 | 643 |  |
| MV2Reverse | Mayfield virus 2 | ACCTGCCGTGCTAACAAATA |  |  |  |
| AHV1Foward | Allermuir Hill virus 1 | TGGGGAAGGAATATTTGCAGGT | T62 | 1498 |  |
| AHV1Reverse | Allermuir Hill virus 1 | GGCATCTTTGAAGATAACCTACGC |  |  |  |
| AHV2Foward | Allermuir Hill virus 2 | TGCTGGTGCTGATGTTACATCT | T62 | 1094 |  |
| AHV2Reverse | Allermuir Hill virus 2 | TTCGAAACACAACTGCAATACA |  |  |  |
| AHV3Foward | Allermuir Hill virus 3 | GGGGCTCGGCTGAATCTAG | PCR 60 | 1283 |  |
| AHV3Reverse | Allermuir Hill virus 3 | TGCAAACAATTAGAGTTGGCCA |  |  |  |
| MLVFoward | Mill Lade virus | TCTCGCAATCCATACGTACTTCA | T62 | 1249 |  |
| MLVReverse | Mill Lade virus | CGTCAACAAGGTCGTTTCTTCC |  |  |  |
| CBVForward | Castleton Burn virus | TTCTCTATCGAGCGGCCTTG | T62 | 850 |  |
| CBVReverse | Castleton Burn virus | TGTGCTTCCATGTAGGCGAA |  |  |  |
| SVForward | Sheriffmuir virus | GTTTTGACCAGCACCAGAGC | PCR 60 | 342 |  |
| SVReverse | Sheriffmuir virus | CCAGTTCGGGGTGGCTAAAT |  |  |  |
| DVForward | Dumyat virus | CGGAGATAACGGAGGTGTGG | T62 | 420 |  |
| DVReverse | Dumyat virus | GCAAGGAGACAAGGCTCCTT |  |  |  |
| CMVForward | Cnoc Mor virus | TCAGCCGAATTAGAATGTGTACA | T62 | 1104 |  |
| CMVReverse | Cnoc Mor virus | GCGCTTTCGAATAGATGCCT |  |  |  |
| ARVForward | Agassiz Rock virus | TGAATGGTAGGAGCATGCGT | PCR 60 | 1350 |  |
| ARVReverse | Agassiz Rock virus | TGTAGTAGATGCCTGGGTTTGA |  |  |  |
| ELVForward | Elf Loch virus | GAACAAGCGCGAGTGGAAAC | PCR 62 | 1337 |  |
| ELVReverse | Elf Loch virus | TCGAGATTATCTGCGTGGCC |  |  |  |
| CCVForward | Clamshell Cave virus | GGCCTCAAGGTATGTTGAATAACA | T62 | 467 |  |
| CCVReverse | Clamshell Cave virus | TGCACTTATTATCTGCTGTCTTAGA |  |  |  |
| BBVForward | Boghill Burn virus | TGGATCACCTGATGGATTCCT | T62 | 1383 |  |
| BBVReverse | Boghill Burn virus | TCGATCTCTCTGTGAGTCTCTGT |  |  |  |
| BHVForward | Black Hill virus | ACATCTATGTGCTGCAGCGA | T62 | 789 |  |
| BHVReverse | Black Hill virus | CCATGACCGTGTGCTAGCAT |  |  |  |
| RLVForward | River Liunaeg virus | ACCAGGTGGAACTCGTGTTT | PCR 60 | 883 |  |
| RLVReverse | River Liunaeg virus | GTACTCTGGACCTTTGCCGT |  |  |  |
| LMVForward | Loch Morlich virus | AGTGGTGGAGATGGAGACGA | PCR 60 | 1479 |  |
| LMVReverse | Loch Morlich virus | CCACAGATACCAGTGGCGTA |  |  |  |
| GVForward | Gorebridge virus | GGATAGATACACTAAAGGATGCTAAAA | T62 | 896 |  |
| GVReverse | Gorebridge virus | ATTCGGTGCATCAAGGAGCA |  |  |  |
| HPLV34Forward | Hubei partiti-like virus 34 | TGGCTTAGATTTAATGCTACGAT | T62 | 520 |  |
| HPLV34Reverse | Hubei partiti-like virus 34 | CCTCATAGCTCCACCAGTAACC |  |  |  |
| SBPV 774F | Slow bee paralysis virus | GAGATGGATMGRCCTGAAGG | T62 |  | Lena Wilfert (*pers comm*) |
| SBPV 1698R | Slow bee paralysis virus | CATGAGCCAKGARTGTGAA |  |  |  |
| ABPV 5088F | Acute bee paralysis virus | CYATGGACACACCCTATGTG | T62 |  | Lena Wilfert (*pers comm*) |
| ABPV 6122R | Acute bee paralysis virus | CGCCATTTTGCTACTTCTCC |  |  |  |

Supplementary Table 4 The genbank IDs of the sequences used to build the host tree. Species added for better estimation of the tree and not used further are shown in bold.

|  | 16S ribosomal RNA gene | Cytochrome oxidase subunit I (COI) gene | Phosphoenolpyruvate carboxykinase | Long-wavelength rhodopsin | Elongation factor-1 alpha gene | Arginine kinase gene |
| --- | --- | --- | --- | --- | --- | --- |
| *Bombus terrestris* | DQ788118.1 | AY181170.1 | EF050865.1 | AF493022.1 | DQ788288.1 | AF492888.1 |
| *Bombus cryptarum* | DQ787995.1 | AY181100.1 | EF050855.1 | AY739461.1 | DQ788175.1 | DQ788416.1 |
| *Bombus lucorum* | DQ788051.1 | AY181120.1 | EF050857.1 | AF493021.1 | DQ788225.1 | AF492887.1 |
| ***Bombus patagiatus*** | DQ788078.1 | AF279499.1 | EF050862.1 | AF493020.1 | DQ788252.1 | AF492886.1 |
| *Bombus lapidarius* | DQ788045.1 | AY181115.1 | EF050902.1 | AF493005.1 | DQ788219.1 | AF492871.1 |
| *Bombus monticola* | DQ788064.1 | AY181132.1 | EF050848.1 | AY739483.1 | DQ788238.1 | DQ788466.1 |
| *Bombus pascuorum* | DQ788077.1 | AY181137.1 | EF050932.1 | AF493001.1 | DQ788251.1 | AF492867.1 |
| *Bombus hortorum* | DQ788024.1 | AY181105.1 | EF050999.1 | AF492987.1 | DQ788200.1 | AF492853.1 |
| *Bombus pratorum* | DQ788087.1 | AY181149.1 | EF050819.1 | AF493033.1 | AF492966.1 | AF492899.1 |
| *Bombus jonellus* | DQ788039.1 | AY181113.1 | EF050814.1 | AY739473.1 | DQ788214.1 | DQ788446.1 |
| *Bombus bohemicus* | DQ787980.1 | AY181181.1 | EF050979.1 | AF492992.1 | AF492925.1 | AF492858.1 |
| ***Bombus waltoni*** | KX791789.1 | KX791763.1 | KX791713.1 | KX791687.1 | KX791663.1 | KX791737.1 |
| ***Apis mellifera*** | L06178.1 | L06178.1 |  |  |  |  |

Supplementary Table 4 The genbank IDs of the sequences used to build the virus tree. Species added for better estimation of the tree and not used further are shown in bold. Viruses excluded from the tree containing only the dsRNA and +ve sense RNA viruses are marked with a cross.

| **Virus** | **Genbank ID** | **Negative Sense?** |
| --- | --- | --- |
| Acute bee paralysis virus | NP_066241.1 |  |
| Allermuir Hill virus 1 | This study |  |
| Allermuir Hill virus 2 | This study |  |
| Allermuir Hill virus 3 | This study |  |
| Aphid lethal paralysis virus | JX045858.1 |  |
| **Beihai paphia shell virus 2** | NC_032494.1 |  |
| **Beihai sobemo-like virus 8** | NC_032972.1 |  |
| **Beihai weivirus-like virus 17** | YP_009336943.1 |  |
| **Beihai zhaovirus-like virus 4** | YP_009337445.1 |  |
| **Big Sioux River virus** | JF423196.1 |  |
| **Black queen cell virus** | NC_003784.1 |  |
| Boghill Burn virus | This study |  |
| **Bovine astrovirus strain BAstV-GX7/CHN/2014** | NC_024297.1 |  |
| Castleton Burn virus | This study |  |
| **Changjiang picorna-like virus 17** | APG79015.1 |  |
| **Chronic bee paralysis virus segment** | KY937971.1 |  |
| Clamshell Cave virus | This study | x |
| **Cricket paralysis virus** | NC_003924.1 |  |
| **Cryphonectria hypovirus 3** | AF188515.1 |  |
| **Deformed wing virus A** | JQ413340.1 |  |
| **Deformed wing virus B** | AY251269.2 |  |
| **Deformed wing virus C** | Unpublished, Lena Wilfert (*pers comm*) |  |
| **Drosophila x virus** | NC_004169.1 |  |
| Dumyat virus | This study |  |
| Elf Loch virus | This study |  |
| **Ganda bee virus** | APT68154.1 | x |
| Gorebridge virus | This study |  |
| **Helminthosporium victoriae virus** | NC_003607.2 |  |
| **Hubei astro-like virus** | NC_032203.1 |  |
| **Hubei chuvirus-like virus 3** | NC_033015 |  |
| **Hubei odonate virus 13** | APG78254.1 |  |
| Hubei partiti-like virus 34 | APG78322.1 |  |
| **Hubei picorna-like virus 15** | APG77985.1 |  |
| **Hubei picorna-like virus 57** | APG78030.1 |  |
| **Hubei tombus-like virus 29** | YP_009336934.1 |  |
| **Iflavirus sp. isolate VDV-2** | KX578271.1 |  |
| **Infectious pancreatic necrosis virus** | NC_001916.1 |  |
| **Israeli acute paralysis virus** | KX421583.1 |  |
| **Kashmir bee virus** | NC_004807.1 |  |
| **Lake Sinai virus 1** | HQ871931.2 |  |
| **Lake Sinai virus 2** | HQ888865.2 |  |
| **Lake Sinai virus 6** | KR021357.1 |  |
| Loch Morlich virus | This study |  |
| Mayfield virus 1 | This study |  |
| Mayfield virus 2 | This study |  |
| Mill Lade virus | This study |  |
| Muthill virus | AMO03223.1 |  |
| **Nodamura virus** | AF174533.1 |  |
| **Norwalk virus** | M87661.2 |  |
| River Liunaeg virus | This study |  |
| **Rubella virus** | NC_001545.2 |  |
| **Sacbrood virus** | NP_049374.1 |  |
| Slow bee paralysis virus | YP_003622540.1 |  |
| **Scaldis River bee virus** | KY053857.1 | x |
| **Seattle Prectang virus** | AOF41423.1 | x |
| Sheriffmuir virus | This study |  |
| **Sindbis virus** | NC_001547.1 |  |
| **Tobacco ringspot virus isolate SK** | KJ556849.1 |  |
| **Varroa destructor virus 3** | KX578272.1 |  |
| **Wenzhou shrimp virus 10** | NC_033164.1 |  |
| **Wenzhou shrimp virus 9** | NC_033300.1 |  |
| **Wuchang Cockroach Virus 1** | YP_009304995.1 | x |
| **Wuhan cricket virus 2** | NC_033764.1 |  |
| **Wuhan insect virus 14** | NC_033473.1 |  |
| **Wuhan spirurian nematodes virus 1** | KX884291.1 |  |
| **Xinzhou nematode virus 1** | YP_009345041.1 |  |

Supplementary Table 6 The correlations from the Brownian motion evolution model between viruses and their 90% shortest posterior intervals

| **Virus 1** | **Virus 2** | **Posterior Mean Correlation** | **Lower 5th Percentile Correlation** | **Upper 95th Percentile Correlation** |
| --- | --- | --- | --- | --- |
| Acute bee paralysis virus | Acute bee paralysis virus | 1.00 | 1.00 | 1.00 |
| Acute bee paralysis virus | Mayfield virus 2 | 0.46 | 0.38 | 0.54 |
| Acute bee paralysis virus | Boghill Burn virus | 0.45 | 0.34 | 0.53 |
| Acute bee paralysis virus | River Luinaeg virus | 0.58 | 0.52 | 0.64 |
| Acute bee paralysis virus | Elf Loch virus | 0.19 | 0.06 | 0.31 |
| Acute bee paralysis virus | Allermuir Hill virus 1 | 0.00 | 0.00 | 0.02 |
| Acute bee paralysis virus | Allermuir Hill virus 2 | 0.00 | 0.00 | 0.02 |
| Acute bee paralysis virus | Allermuir Hill virus 3 | 0.00 | 0.00 | 0.02 |
| Acute bee paralysis virus | Hubei partiti-like virus 34 | 0.15 | 0.05 | 0.26 |
| Acute bee paralysis virus | Castleton Burn virus | 0.01 | 0.00 | 0.05 |
| Acute bee paralysis virus | Gorebridge virus | 0.69 | 0.64 | 0.73 |
| Acute bee paralysis virus | Loch Morlich virus | 0.58 | 0.52 | 0.64 |
| Acute bee paralysis virus | Mayfield virus 1 | 0.46 | 0.38 | 0.54 |
| Acute bee paralysis virus | Mill Lade virus | 0.00 | 0.00 | 0.02 |
| Acute bee paralysis virus | Slow bee paralysis virus | 0.58 | 0.52 | 0.64 |
| Acute bee paralysis virus | Dumyat virus | 0.19 | 0.07 | 0.29 |
| Acute bee paralysis virus | Sheriffmuir virus | 0.19 | 0.07 | 0.29 |
| Acute bee paralysis virus | Clamshell Cave virus | 0.17 | 0.02 | 0.29 |
| Mayfield virus 2 | Mayfield virus 2 | 1.00 | 1.00 | 1.00 |
| Mayfield virus 2 | Boghill Burn virus | 0.51 | 0.36 | 0.62 |
| Mayfield virus 2 | River Luinaeg virus | 0.46 | 0.38 | 0.54 |
| Mayfield virus 2 | Elf Loch virus | 0.19 | 0.06 | 0.31 |
| Mayfield virus 2 | Allermuir Hill virus 1 | 0.00 | 0.00 | 0.02 |
| Mayfield virus 2 | Allermuir Hill virus 2 | 0.00 | 0.00 | 0.02 |
| Mayfield virus 2 | Allermuir Hill virus 3 | 0.00 | 0.00 | 0.02 |
| Mayfield virus 2 | Hubei partiti-like virus 34 | 0.15 | 0.05 | 0.26 |
| Mayfield virus 2 | Castleton Burn virus | 0.01 | 0.00 | 0.05 |
| Mayfield virus 2 | Gorebridge virus | 0.46 | 0.38 | 0.54 |
| Mayfield virus 2 | Loch Morlich virus | 0.46 | 0.38 | 0.54 |
| Mayfield virus 2 | Mayfield virus 1 | 0.99 | 0.99 | 1.00 |
| Mayfield virus 2 | Mill Lade virus | 0.00 | 0.00 | 0.02 |
| Mayfield virus 2 | Slow bee paralysis virus | 0.46 | 0.38 | 0.54 |
| Mayfield virus 2 | Dumyat virus | 0.19 | 0.07 | 0.29 |
| Mayfield virus 2 | Sheriffmuir virus | 0.19 | 0.07 | 0.29 |
| Mayfield virus 2 | Clamshell Cave virus | 0.17 | 0.02 | 0.29 |
| Boghill Burn virus | Boghill Burn virus | 1.00 | 1.00 | 1.00 |
| Boghill Burn virus | River Luinaeg virus | 0.45 | 0.34 | 0.53 |
| Boghill Burn virus | Elf Loch virus | 0.19 | 0.06 | 0.31 |
| Boghill Burn virus | Allermuir Hill virus 1 | 0.00 | 0.00 | 0.02 |
| Boghill Burn virus | Allermuir Hill virus 2 | 0.00 | 0.00 | 0.02 |
| Boghill Burn virus | Allermuir Hill virus 3 | 0.00 | 0.00 | 0.02 |
| Boghill Burn virus | Hubei partiti-like virus 34 | 0.15 | 0.05 | 0.26 |
| Boghill Burn virus | Castleton Burn virus | 0.01 | 0.00 | 0.05 |
| Boghill Burn virus | Gorebridge virus | 0.45 | 0.34 | 0.53 |
| Boghill Burn virus | Loch Morlich virus | 0.45 | 0.34 | 0.53 |
| Boghill Burn virus | Mayfield virus 1 | 0.51 | 0.36 | 0.62 |
| Boghill Burn virus | Mill Lade virus | 0.00 | 0.00 | 0.02 |
| Boghill Burn virus | Slow bee paralysis virus | 0.45 | 0.34 | 0.53 |
| Boghill Burn virus | Dumyat virus | 0.19 | 0.07 | 0.29 |
| Boghill Burn virus | Sheriffmuir virus | 0.19 | 0.07 | 0.29 |
| Boghill Burn virus | Clamshell Cave virus | 0.17 | 0.02 | 0.29 |
| River Luinaeg virus | River Luinaeg virus | 1.00 | 1.00 | 1.00 |
| River Luinaeg virus | Elf Loch virus | 0.19 | 0.06 | 0.31 |
| River Luinaeg virus | Allermuir Hill virus 1 | 0.00 | 0.00 | 0.02 |
| River Luinaeg virus | Allermuir Hill virus 2 | 0.00 | 0.00 | 0.02 |
| River Luinaeg virus | Allermuir Hill virus 3 | 0.00 | 0.00 | 0.02 |
| River Luinaeg virus | Hubei partiti-like virus 34 | 0.15 | 0.05 | 0.26 |
| River Luinaeg virus | Castleton Burn virus | 0.01 | 0.00 | 0.05 |
| River Luinaeg virus | Gorebridge virus | 0.58 | 0.52 | 0.64 |
| River Luinaeg virus | Loch Morlich virus | 0.67 | 0.62 | 0.71 |
| River Luinaeg virus | Mayfield virus 1 | 0.46 | 0.38 | 0.54 |
| River Luinaeg virus | Mill Lade virus | 0.00 | 0.00 | 0.02 |
| River Luinaeg virus | Slow bee paralysis virus | 0.67 | 0.62 | 0.72 |
| River Luinaeg virus | Dumyat virus | 0.19 | 0.07 | 0.29 |
| River Luinaeg virus | Sheriffmuir virus | 0.19 | 0.07 | 0.29 |
| River Luinaeg virus | Clamshell Cave virus | 0.17 | 0.02 | 0.29 |
| Elf Loch virus | Elf Loch virus | 1.00 | 1.00 | 1.00 |
| Elf Loch virus | Allermuir Hill virus 1 | 0.00 | 0.00 | 0.00 |
| Elf Loch virus | Allermuir Hill virus 2 | 0.00 | 0.00 | 0.00 |
| Elf Loch virus | Allermuir Hill virus 3 | 0.00 | 0.00 | 0.00 |
| Elf Loch virus | Hubei partiti-like virus 34 | 0.14 | 0.03 | 0.25 |
| Elf Loch virus | Castleton Burn virus | 0.00 | 0.00 | 0.03 |
| Elf Loch virus | Gorebridge virus | 0.19 | 0.06 | 0.31 |
| Elf Loch virus | Loch Morlich virus | 0.19 | 0.06 | 0.31 |
| Elf Loch virus | Mayfield virus 1 | 0.19 | 0.06 | 0.31 |
| Elf Loch virus | Mill Lade virus | 0.00 | 0.00 | 0.00 |
| Elf Loch virus | Slow bee paralysis virus | 0.19 | 0.06 | 0.31 |
| Elf Loch virus | Dumyat virus | 0.25 | 0.07 | 0.41 |
| Elf Loch virus | Sheriffmuir virus | 0.25 | 0.07 | 0.41 |
| Elf Loch virus | Clamshell Cave virus | 0.17 | 0.00 | 0.32 |
| Allermuir Hill virus 1 | Allermuir Hill virus 1 | 1.00 | 1.00 | 1.00 |
| Allermuir Hill virus 1 | Allermuir Hill virus 2 | 0.98 | 0.97 | 0.99 |
| Allermuir Hill virus 1 | Allermuir Hill virus 3 | 0.98 | 0.97 | 0.99 |
| Allermuir Hill virus 1 | Hubei partiti-like virus 34 | 0.00 | 0.00 | 0.02 |
| Allermuir Hill virus 1 | Castleton Burn virus | 0.15 | 0.03 | 0.27 |
| Allermuir Hill virus 1 | Gorebridge virus | 0.00 | 0.00 | 0.02 |
| Allermuir Hill virus 1 | Loch Morlich virus | 0.00 | 0.00 | 0.02 |
| Allermuir Hill virus 1 | Mayfield virus 1 | 0.00 | 0.00 | 0.02 |
| Allermuir Hill virus 1 | Mill Lade virus | 0.74 | 0.68 | 0.78 |
| Allermuir Hill virus 1 | Slow bee paralysis virus | 0.00 | 0.00 | 0.02 |
| Allermuir Hill virus 1 | Dumyat virus | 0.00 | 0.00 | 0.02 |
| Allermuir Hill virus 1 | Sheriffmuir virus | 0.00 | 0.00 | 0.02 |
| Allermuir Hill virus 1 | Clamshell Cave virus | 0.00 | 0.00 | 0.00 |
| Allermuir Hill virus 2 | Allermuir Hill virus 2 | 1.00 | 1.00 | 1.00 |
| Allermuir Hill virus 2 | Allermuir Hill virus 3 | 0.98 | 0.97 | 0.99 |
| Allermuir Hill virus 2 | Hubei partiti-like virus 34 | 0.00 | 0.00 | 0.02 |
| Allermuir Hill virus 2 | Castleton Burn virus | 0.15 | 0.03 | 0.27 |
| Allermuir Hill virus 2 | Gorebridge virus | 0.00 | 0.00 | 0.02 |
| Allermuir Hill virus 2 | Loch Morlich virus | 0.00 | 0.00 | 0.02 |
| Allermuir Hill virus 2 | Mayfield virus 1 | 0.00 | 0.00 | 0.02 |
| Allermuir Hill virus 2 | Mill Lade virus | 0.74 | 0.68 | 0.78 |
| Allermuir Hill virus 2 | Slow bee paralysis virus | 0.00 | 0.00 | 0.02 |
| Allermuir Hill virus 2 | Dumyat virus | 0.00 | 0.00 | 0.02 |
| Allermuir Hill virus 2 | Sheriffmuir virus | 0.00 | 0.00 | 0.02 |
| Allermuir Hill virus 2 | Clamshell Cave virus | 0.00 | 0.00 | 0.00 |
| Allermuir Hill virus 3 | Allermuir Hill virus 3 | 1.00 | 1.00 | 1.00 |
| Allermuir Hill virus 3 | Hubei partiti-like virus 34 | 0.00 | 0.00 | 0.02 |
| Allermuir Hill virus 3 | Castleton Burn virus | 0.15 | 0.03 | 0.27 |
| Allermuir Hill virus 3 | Gorebridge virus | 0.00 | 0.00 | 0.02 |
| Allermuir Hill virus 3 | Loch Morlich virus | 0.00 | 0.00 | 0.02 |
| Allermuir Hill virus 3 | Mayfield virus 1 | 0.00 | 0.00 | 0.02 |
| Allermuir Hill virus 3 | Mill Lade virus | 0.74 | 0.68 | 0.78 |
| Allermuir Hill virus 3 | Slow bee paralysis virus | 0.00 | 0.00 | 0.02 |
| Allermuir Hill virus 3 | Dumyat virus | 0.00 | 0.00 | 0.02 |
| Allermuir Hill virus 3 | Sheriffmuir virus | 0.00 | 0.00 | 0.02 |
| Allermuir Hill virus 3 | Clamshell Cave virus | 0.00 | 0.00 | 0.00 |
| Hubei partiti-like virus 34 | Hubei partiti-like virus 34 | 1.00 | 1.00 | 1.00 |
| Hubei partiti-like virus 34 | Castleton Burn virus | 0.01 | 0.00 | 0.05 |
| Hubei partiti-like virus 34 | Gorebridge virus | 0.15 | 0.05 | 0.26 |
| Hubei partiti-like virus 34 | Loch Morlich virus | 0.15 | 0.05 | 0.26 |
| Hubei partiti-like virus 34 | Mayfield virus 1 | 0.15 | 0.05 | 0.26 |
| Hubei partiti-like virus 34 | Mill Lade virus | 0.00 | 0.00 | 0.02 |
| Hubei partiti-like virus 34 | Slow bee paralysis virus | 0.15 | 0.05 | 0.26 |
| Hubei partiti-like virus 34 | Dumyat virus | 0.15 | 0.04 | 0.28 |
| Hubei partiti-like virus 34 | Sheriffmuir virus | 0.15 | 0.04 | 0.28 |
| Hubei partiti-like virus 34 | Clamshell Cave virus | 0.13 | 0.00 | 0.24 |
| Castleton Burn virus | Castleton Burn virus | 1.00 | 1.00 | 1.00 |
| Castleton Burn virus | Gorebridge virus | 0.01 | 0.00 | 0.05 |
| Castleton Burn virus | Loch Morlich virus | 0.01 | 0.00 | 0.05 |
| Castleton Burn virus | Mayfield virus 1 | 0.01 | 0.00 | 0.05 |
| Castleton Burn virus | Mill Lade virus | 0.15 | 0.03 | 0.27 |
| Castleton Burn virus | Slow bee paralysis virus | 0.01 | 0.00 | 0.05 |
| Castleton Burn virus | Dumyat virus | 0.01 | 0.00 | 0.04 |
| Castleton Burn virus | Sheriffmuir virus | 0.01 | 0.00 | 0.04 |
| Castleton Burn virus | Clamshell Cave virus | 0.00 | 0.00 | 0.02 |
| Gorebridge virus | Gorebridge virus | 1.00 | 1.00 | 1.00 |
| Gorebridge virus | Loch Morlich virus | 0.58 | 0.52 | 0.64 |
| Gorebridge virus | Mayfield virus 1 | 0.46 | 0.38 | 0.54 |
| Gorebridge virus | Mill Lade virus | 0.00 | 0.00 | 0.02 |
| Gorebridge virus | Slow bee paralysis virus | 0.58 | 0.52 | 0.64 |
| Gorebridge virus | Dumyat virus | 0.19 | 0.07 | 0.29 |
| Gorebridge virus | Sheriffmuir virus | 0.19 | 0.07 | 0.29 |
| Gorebridge virus | Clamshell Cave virus | 0.17 | 0.02 | 0.29 |
| Loch Morlich virus | Loch Morlich virus | 1.00 | 1.00 | 1.00 |
| Loch Morlich virus | Mayfield virus 1 | 0.46 | 0.38 | 0.54 |
| Loch Morlich virus | Mill Lade virus | 0.00 | 0.00 | 0.02 |
| Loch Morlich virus | Slow bee paralysis virus | 0.73 | 0.68 | 0.77 |
| Loch Morlich virus | Dumyat virus | 0.19 | 0.07 | 0.29 |
| Loch Morlich virus | Sheriffmuir virus | 0.19 | 0.07 | 0.29 |
| Loch Morlich virus | Clamshell Cave virus | 0.17 | 0.02 | 0.29 |
| Mayfield virus 1 | Mayfield virus 1 | 1.00 | 1.00 | 1.00 |
| Mayfield virus 1 | Mill Lade virus | 0.00 | 0.00 | 0.02 |
| Mayfield virus 1 | Slow bee paralysis virus | 0.46 | 0.38 | 0.54 |
| Mayfield virus 1 | Dumyat virus | 0.19 | 0.07 | 0.29 |
| Mayfield virus 1 | Sheriffmuir virus | 0.19 | 0.07 | 0.29 |
| Mayfield virus 1 | Clamshell Cave virus | 0.17 | 0.02 | 0.29 |
| Mill Lade virus | Mill Lade virus | 1.00 | 1.00 | 1.00 |
| Mill Lade virus | Slow bee paralysis virus | 0.00 | 0.00 | 0.02 |
| Mill Lade virus | Dumyat virus | 0.00 | 0.00 | 0.02 |
| Mill Lade virus | Sheriffmuir virus | 0.00 | 0.00 | 0.02 |
| Mill Lade virus | Clamshell Cave virus | 0.00 | 0.00 | 0.00 |
| Slow bee paralysis virus | Slow bee paralysis virus | 1.00 | 1.00 | 1.00 |
| Slow bee paralysis virus | Dumyat virus | 0.19 | 0.07 | 0.29 |
| Slow bee paralysis virus | Sheriffmuir virus | 0.19 | 0.07 | 0.29 |
| Slow bee paralysis virus | Clamshell Cave virus | 0.17 | 0.02 | 0.29 |
| Dumyat virus | Dumyat virus | 1.00 | 1.00 | 1.00 |
| Dumyat virus | Sheriffmuir virus | 0.96 | 0.94 | 0.97 |
| Dumyat virus | Clamshell Cave virus | 0.16 | 0.00 | 0.31 |
| Sheriffmuir virus | Sheriffmuir virus | 1.00 | 1.00 | 1.00 |
| Sheriffmuir virus | Clamshell Cave virus | 0.16 | 0.00 | 0.31 |
| Clamshell Cave virus | Clamshell Cave virus | 1.00 | 1.00 | 1.00 |

**Supplementary Table 7** The putative viral contigs found by the *de novo* assembly of raw RNAseq reads after bioinformatic checking

| **Putative viral contig** | **Clade** | **Genome structure** | **Seque-nce length** | **Closest blastx match** | **Query cover/ Percent identity** | **Valida -ted by PCR?** | **Prevalence data acquired?** | **Notes** |
| --- | --- | --- | --- | --- | --- | --- | --- | --- |
| Densovirus 1 | Parvoviridae | positive sense ssDNA | 811 | Neodiprion lecontei nucleopolyhedrovirus (YP_025282.1) | 39%/ 42% |  |  | One incomplete ORF; predicted protein contains the Parvovirus coat protein VP1 (PF08398) motif. While top hit is to a baculoviridae, all other information implies that densovirus is the correct characterization. |
| Densovirus2 | Parvoviridae | positive sense ssDNA | 1345 | Diaphorina citri densovirus (ALV85426.1) | 67%/ 28% |  |  | One incomplete ORF; no predicted motifs. Blastx hits against RNA helicases. |
| Agassiz Rock virus | Reo | dsRNA | 1385 | Hubei odonate virus 14 (APG79163.1) | 72%/ 40% | ✓ | ✓ | One incomplete ORF; no predicted motifs. Blastx hits against RdRps. |
| Cnoc Mor virus | Reo | dsRNA | 1116 | Grange virus (AMO03252.1) | 80%/ 38% | ✓ | ✓ | One incomplete ORF; no predicted motifs. Blastx hits against RdRps. |
| Reoviridae 1 | Reo | dsRNA | 1048 | Hubei odonate virus 14 (APG79163.1) | 96%/ 47% |  |  | One incomplete ORF; no predicted motifs. Blastx hits against RdRps. Aligns with Elf Loch virus and Agassiz Rock virus, potentially part of Cnoc Mor virus. |
| Elf Loch virus | Reo | dsRNA | 2996 | Hubei odonate virus 14 (APG79163.1) | 87%/ 44% | ✓ | ✓ | One incomplete ORF; no predicted motifs, but manual search found GDD motif of RdRp. Blastx hits against RdRps. |
| Dumyat virus | Toti-Chryso | dsRNA | 4981 | Leptopilina boulardi Toti-like virus (YP_009072448.1) | 46%/ 48% | ✓ | ✓ | One incomplete ORF; predicted protein contains the RdRp 4 (PF02123) motif. |
| Sheriffmuir virus | Toti-Chryso | dsRNA | 549 | Leptopilina boulardi Toti-like virus (YP_009072448.1) | 100%/ 63% | ✓ | ✓ | One incomplete ORF; no predicted motifs, but manual search found GDD motif of RdRp. Blastx hits against RdRps. |
| Clamshell Cave virus | Bunya-Arena | negative sense ssRNA | 505 | Ganda bee virus (APT68154.1) | 99%/ 86% | ✓ | ✓ | One incomplete ORF; predicted protein contains the Bunyavirus RdRp (PF04196) motif. |
| Bunyaviridae 1 | Bunya-Arena | negative sense ssRNA | 736 | Ganda bee virus (APT68155.1) | 81%/ 60% |  |  | One incomplete ORF; no predicted motifs. Hit against Bunyaviridae glycoprotein. Potentially Clamshell Cave virus M segment. |
| Bunyaviridae 2 | Bunya-Arena | negative sense ssRNA | 747 | Ganda bee virus (APT68156.1) | 99%/ 84% |  |  | One incomplete ORF; no predicted motifs. Hit against Bunyaviridae nucleoprotein. Potentially Clamshell Cave virus S segment. |
| Phlebovirus 1 | Bunya-Arena | negative sense ssRNA | 2188 | EgAN 1825-61 virus (AEL29653.1) | 61%/ 25% |  |  | One incomplete ORF; predicted protein contains Phlebovirus glycoprotein G2 (PF07245). |
| Orthomyxovirus 1 | Orthomyxovidae | negative sense ssRNA | 1449 | Hubei orthoptera virus 6 (APG77906.1) | 90%/ 26% |  |  | One incomplete ORF; predicted protein contains the Influenza RdRp subunit PB1 (PF00602) motif. |
| Allermuir Hill virus 1 | Hepe-Virga | positive sense ssRNA | 7586 | Xinzhou nematode virus 1 (YP_009345041.1) | 39%/ 34% | ✓ | ✓ | Two putative ORFs; one containing the FtsJ-like methyltransferase (PF01728), RdRp 1 (PF00978) and viral (Superfamily 1) RNA helicase (PF01443) motifs and one with no predicted motifs and no blast hits. |
| Allermuir Hill virus 2 | Hepe-Virga | positive sense ssRNA | 9078 | Xinzhou nematode virus 1 (YP_009345041.1) | 49%/ 34% | ✓ | ✓ | Two putative ORFs; one containing the FtsJ-like methyltransferase (PF01728), viral methyltransferase (PF01660), RdRp 1 (PF00978) and viral (Superfamily 1) RNA helicase (PF01443) motifs and one with no one with no predicted motifs and no blast hits. |
| Allermuir Hill virus 3 | Hepe-Virga | positive sense ssRNA | 6339 | Xinzhou nematode virus 1 (YP_009345041.1) | 47%/ 34% | ✓ | ✓ | Three putative ORFs; one containing the RdRp 1 (PF00978) and viral (Superfamily 1) RNA helicase (PF01443) motifs, one with no one with no predicted motifs and no blast hits, and one with no predicted motifs and blast hits against hypothetical viral proteins. |
| Mill Lade virus | Hepe-Virga | positive sense ssRNA | 3152 | Aedes camptorhynchus negev-like virus (YP_009388601.1) | 44%/ 64% | ✓ | ✓ | One incomplete ORF; predicted protein contains the RdRp 2 (PF00978) and viral (Superfamily 1) RNA helicase (PF01443) motifs. |
| Virga-like virus 1 | Hepe-Virga | positive sense ssRNA | 1681 | Blackford virus (AMO03220.1) | 63%/ 63% |  |  | One incomplete ORF; predicted protein contains the RdRp 2 (PF00978) motif. |
| Virga-like virus 2 | Hepe-Virga | positive sense ssRNA | 962 | Hubei virga-like virus 2 (APG77663.1) | 92%/ 31% |  |  | One incomplete ORF; predicted protein contains the viral (Superfamily 1) RNA helicase (PF01443) and AAA (PF13245) motifs. |
| Virga-like virus 3 | Hepe-Virga | positive sense ssRNA | 2265 | Aedes camptorhynchus negev-like virus (YP_009388603.1) | 28%/ 39% |  |  | Multiple short ORFs; no predicted motifs, blast hits against hypothetical viral proteins. |
| Virga-like virus 4 | Hepe-Virga | positive sense ssRNA | 1840 | Hubei virga-like virus 17 (YP_009337715.1) | 52%/ 34% |  |  | One incomplete ORF; predicted protein contains the viral methyltransferase (PF01660) motif. |
| Black Hill virus | Picorna-Calici | positive sense ssRNA | 1332 | Sacbrood virus (AID58097.1) | 92%/ 55% | ✓ |  | One incomplete ORF; predicted protein contains the RdRp 1 (PF00680) motif. |
| Boghill Burn virus | Picorna-Calici | positive sense ssRNA | 9873 | Hubei picorna-like virus 57 (APG78030.1) | 60%/ 98% | ✓ | ✓ | Two putative ORFs; one containing predicted the RNA helicase (PF00680) and RdRp 1 (PF00910) motifs, one with no predicted motifs. The closest well-studied virus is Acyrthosiphon pisum virus, which contains a second ORF at the 3’ end of its first with no start codon. This ORF is translated by -1 translational frameshift during protein synthesis (van der Wilk, Dullemans, Verbeek, & Van den Heuvel, 1997). This contig contains similar shift from the protein coding sequence lying in the +3 frame (as defined from the beginning) into the +2 frame, with partial overlap of the ORFs. |
| Gorebridge virus | Picorna-Calici | positive sense ssRNA | 5639 | Hubei picorna-like virus 15 (APG77985.1) | 45%/ 97% | ✓ | ✓ | Two putative ORFs; one containing the RdRp 1 (PF00680) motif and one containing the CRPV capsid protein like (PF08762) and picornavirus capsid protein (PF00073) motifs. Apparent standard Dicistroviridae organisation. |
| Picornavirales 1 | Picorna-Calici | positive sense ssRNA | 1819 | Hubei picorna-like virus 15 (APG77986.1) | 75%/ 90% |  |  | One incomplete ORF; predicted protein contains the CRPV capsid protein like (PF08762) and picornavirus capsid protein (PF00073) motifs. |
| Loch Morlich virus | Picorna-Calici | positive sense ssRNA | 3817 | Sacbrood virus (AJA38040.1) | 60%/ 40% | ✓ | ✓ | One incomplete ORF; predicted protein contains the RdRp 1 (PF00680) motif. |
| Mayfield virus 1 | Picorna-Calici | positive sense ssRNA | 8948 | Wenzhou picorna-like virus 47 (APG78496.1) | 53%/ 29% | ✓ | ✓ | One complete ORF; predicted protein contains the RdRp 1 (PF00680) and RNA helicase (PF00910) motif. |
| Mayfield virus 2 | Picorna-Calici | positive sense ssRNA | 9019 | Wenzhou picorna-like virus 47 (APG78496.1) | 53%/ 29% | ✓ | ✓ | One complete ORF; predicted protein contains the RdRp 1 (PF00680) and RNA helicase (PF00910) motif. |
| Nepovirus 1 | Picorna-Calici | positive sense ssRNA | 2426 | Beet ringspot virus (X04062.1) | 89%/ 95% |  |  | One incomplete ORF; predicted protein contains the Nepovirus coat protein, N-terminal domain (PF03689), Nepovirus coat protein, central domain (PF03391) and Nepovirus coat protein, C-terminal domain (PF03688) motifs. |
| Nepovirus 2 | Picorna-Calici | positive sense ssRNA | 1282 | Soybean latent spherical virus (YP_009330271.1) | 58%/ 47% |  |  | One incomplete ORF; no predicted motifs. Blastx hits against RdRps. |
| Nepovirus 3 | Picorna-Calici | positive sense ssRNA | 3816 | Tomato black ring virus (CAA56792.1) | 91%/ 60% |  |  | One incomplete ORF; predicted protein contains the Nepovirus coat protein, N-terminal domain (PF03689), Nepovirus coat protein, central domain (PF03391) and Nepovirus coat protein, C-terminal domain (PF03688) motifs. |
| Picornavirales 2 | Picorna-Calici | positive sense ssRNA | 1798 | Kilifi virus (YP_009140560.1) | 19%/ 30% |  |  | One incomplete ORF and one complete ORF; no predicted motifs. No blastx hits against identifed proteins. |
| River Liunaeg virus | Picorna-Calici | positive sense ssRNA | 1955 | Hubei tick virus 2 (YP_009336542.1) | 91%/ 40% | ✓ | ✓ | One incomplete ORF; predicted protein contains the RdRp 1 (PF00680) motif. |
| Castleton Burn virus | Tombus-Noda | positive sense ssRNA | 2714 | Shangao tombus-like virus 1 (APG76298.1) | 44%/ 43% | ✓ | ✓ | One incomplete ORF and one complete ORF; predicted protein contains the RdRp 3 (PF00998) motif and one ORF contains no predicted motifs. No blastx hits against the ORF with no predicted motifs. |
| Nodavirus 1 | Tombus-Noda | positive sense ssRNA | 1011 | Wuhan nodavirus (ABB71128.1) | 78%/ 66% |  |  | One incomplete ORF; predicted protein contains the nodavirus capsid protein (PF11729) motif. |

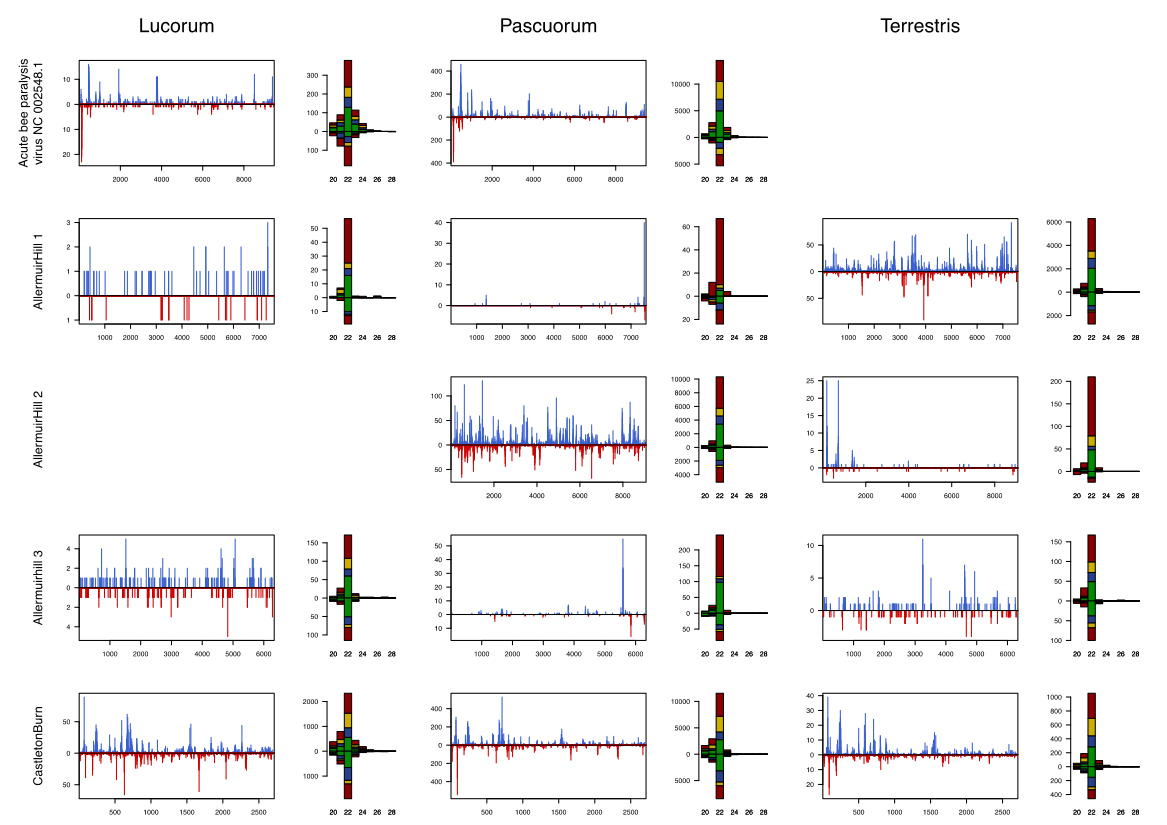

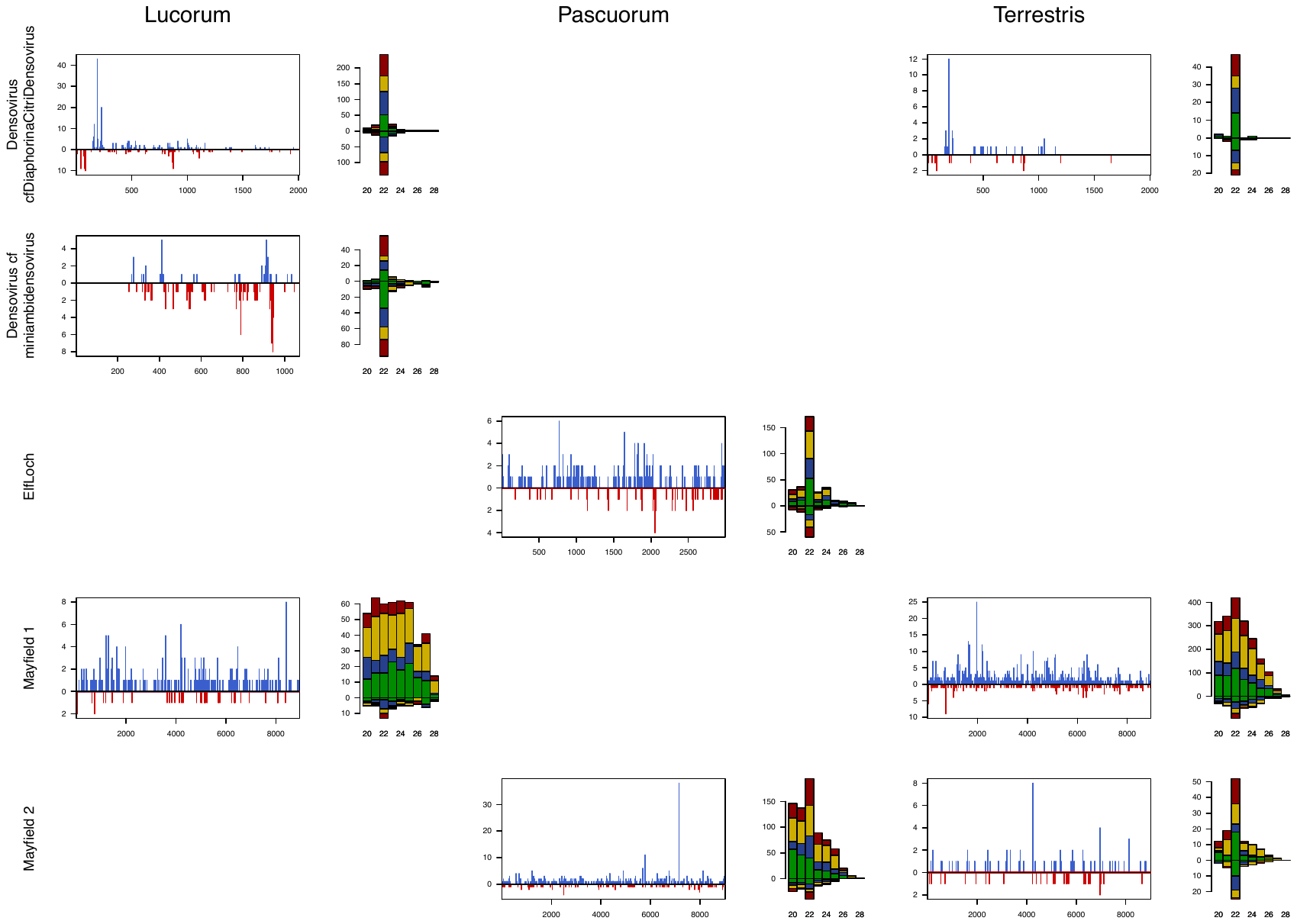

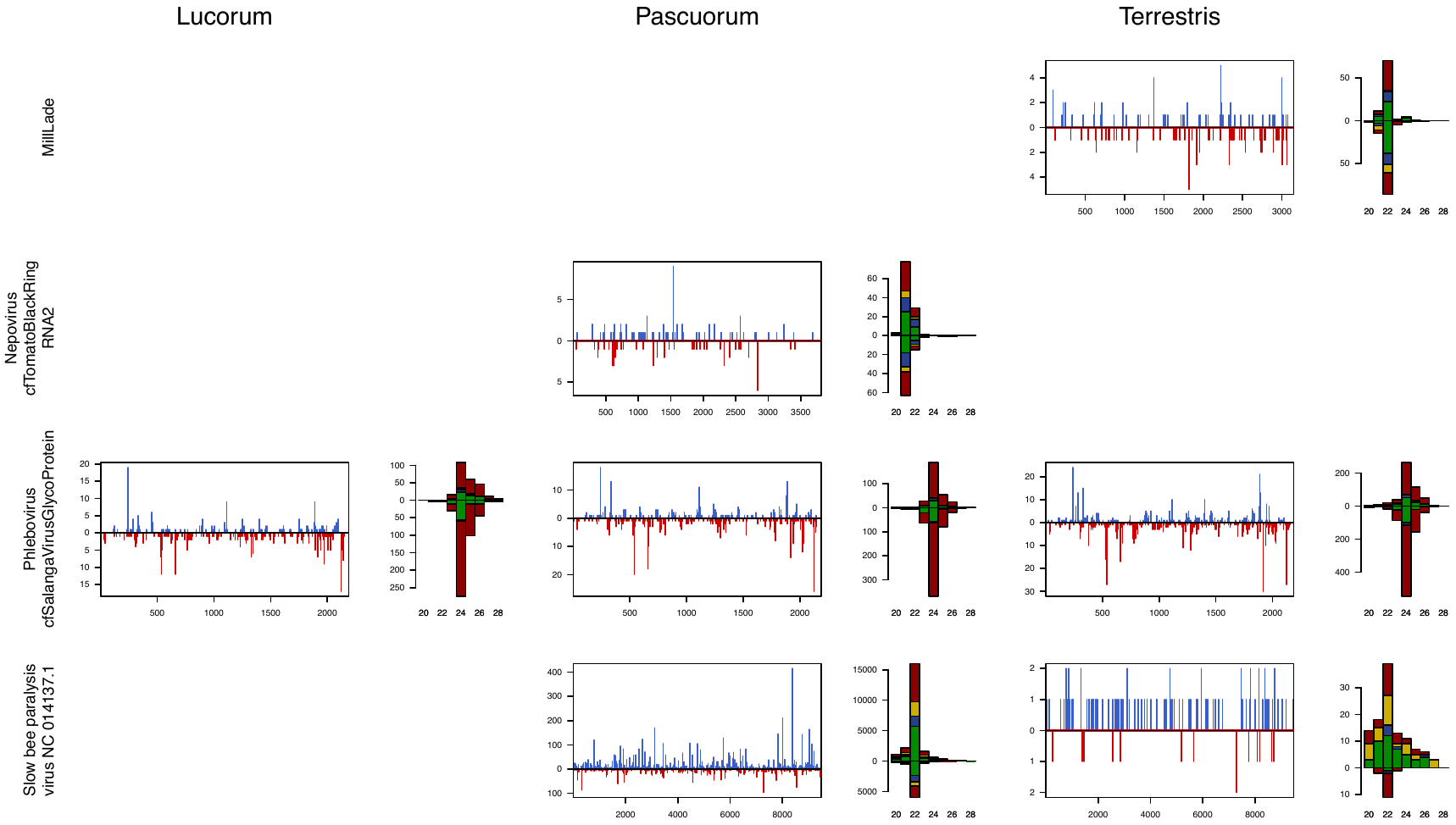

**Supplementary Figure 1** The mapping of small RNA reads to each virus with at least 50 mapping reads, and their read size spectra. Blue lines represent reads mapping to the forward strand at that genomic position, red lines represent reads mapping to the reverse strand. The histogram of read size spectra shows the count of reads of each length mapping in the positive (above) and negative (below) directions. The colouring of each bar shows the counts of the reads beginning with each 5’ base (red-U, blue-C, green-A, yellow-G).

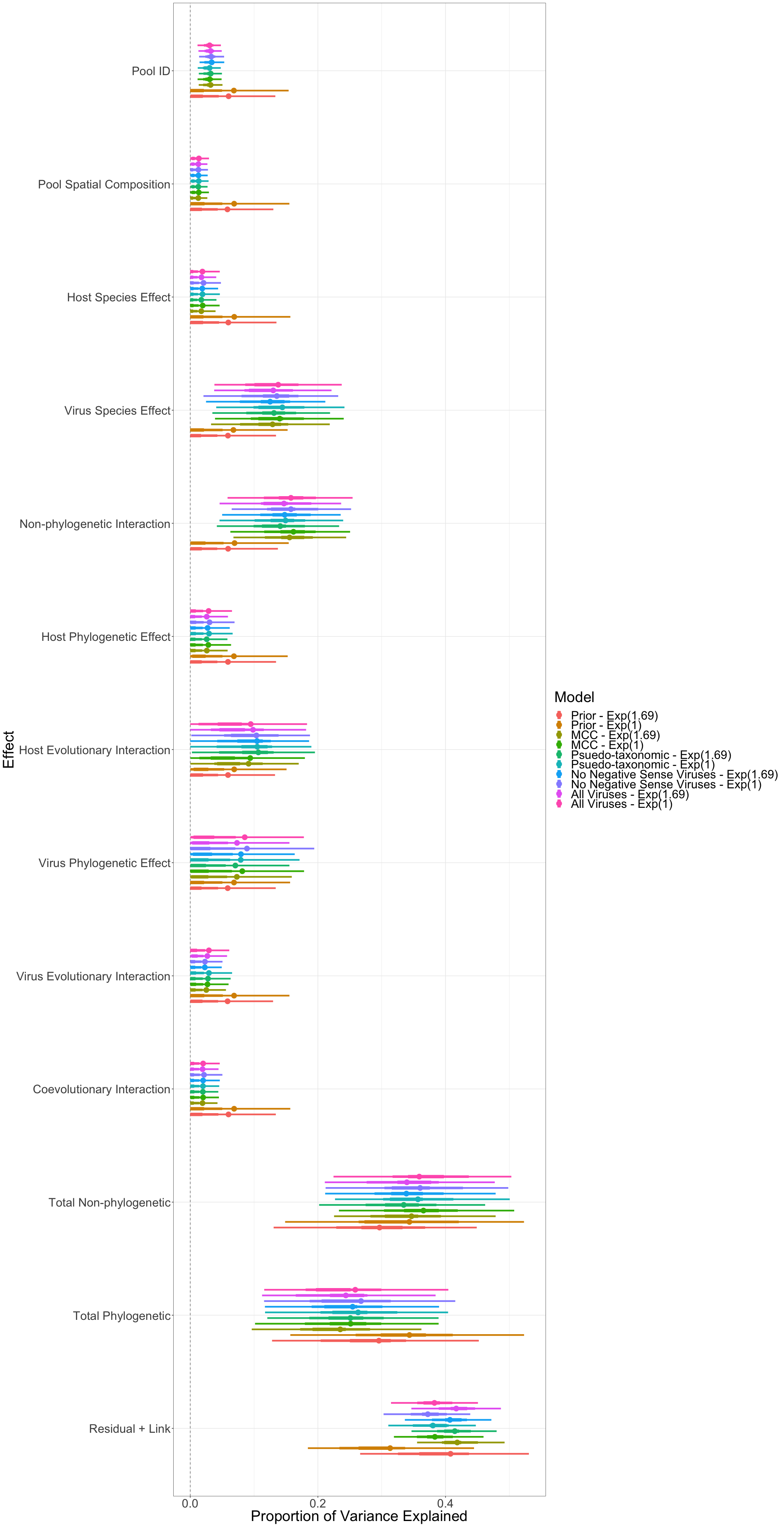

**Supplementary Figure 2** Comparison of estimated proportion of variance in prevalence explained by different parameters between models showing consistency of estimates under different priors. As each factor explains a proportion of the attributed variance in the model total over all factors must sum to 1. For each parameter, the circle represents the modal estimate, the thick bars represent the 50% shortest posterior density interval and the thin bars represent the 90% shortest posterior density interval. “Pool ID” is the proportion of the total variation in prevalence explained by pools within species differing in the degree to which they were infected by viruses. “Spatial Composition” is the proportion of the total variation in prevalence explained by the combination of locations from which the bees in the pool originate. “Host Species Effect” is the proportion of variation in prevalence explained by hosts having different average viral prevalences. “Virus Species Effect” is the proportion of variation in prevalence explained by viruses differing in their average prevalences. “Non-phylogenetic Interaction” is the proportion of variation in prevalence explained by host-virus combinations differing in the their average prevalences beyond that which would be expected by their host and virus species effects alone. “Host Phylogenetic Effect” is the proportion of variation in prevalence explained by hosts having average viral prevalences correlated across the host phylogeny. “Virus Phylogenetic Effect” is the proportion of variation in prevalence explained by viruses having average prevalences correlated across the viral phylogeny. “Host Evolutionary Interaction” is the proportion of variation in prevalence explained by related hosts having correlated viral assemblages. “Virus Evolutionary Interaction” is the proportion of the variation explained by related viruses having correlated host assemblages. “Coevolutionary Interaction” is the proportion of the variation explained by related hosts having similar prevalences of related viruses. “Total Non-phylogenetic” is the proportion of the variation that can be explained by terms not involving the host and virus phylogeny and excluding the residual (“Host Species Effect”, “Virus Species Effect”, “Pool ID”, “Spatial Composition Effect”, “Non-phylogenetic Interaction”). “Total phylogenetic” is the proportion of the variation that can be explained by terms involving a host or virus phylogeny (“Host Phylogenetic Effect”, “Virus Phylogenetic Effect”, “Host Evolutionary Interaction”, “Virus Evolutionary Interaction”, “Coevolutionary Interaction”). “Residual + Link” is the proportion of the total variance that is explained by the residual variance and variance of the logistic distribution (π^2^/3).
